# Passing the quadrat: inferring biodiversity change over time and across investigators

**DOI:** 10.1101/2024.11.05.622130

**Authors:** Robin Elahi, Fiorenza Micheli, James Watanabe

## Abstract

Long-term ecological monitoring inevitably requires a ‘passing of the quadrat’ from one investigator to another. Here we present the challenge and opportunity of inferring biodiversity change over time and across investigators using a rocky intertidal case study. The intertidal transect was surveyed initially by Willis Hewatt in 1931-1933, then resurveyed in detail by Rafe Sagarin, Sarah Gilman, and their mentors in 1993. In total, the transect has been resurveyed 17 times over the past three decades, with annual surveys nested within five distinct investigator eras. We addressed two goals with our dataset. First, we asked whether temporal change in several biodiversity metrics (species richness, Hill-Shannon and Hill-Simpson diversity, spatial beta diversity) was sensitive to investigator era. Second, we identified recent ecological ‘winners’ and ‘losers’, relative to the historical baseline, and asked whether such designations can change over time (1993-2023). We show that investigators contributed to variable trends in species richness over time, and we hypothesize that sampling effort contributed to this effect. In contrast, we highlight declines in the biodiversity of common species (Hill-Shannon) and dominant species (Hill-Simpson) over time, in the absence of investigator effects. These declines were associated with the rise of a few numerically dominant species and a trend of increasing spatial similarity. Relative to the historical baseline, ecological winners included certain barnacles, anemones, and turban snails, while losers included mussels, urchins, limpets and a host of more cryptic taxa. However, we show that such designations can be complicated by non-linear population trajectories and discuss mechanisms related to climate change versus potential artifacts related to investigator effort. We finish with recommendations and emphasize the value of marine stations in providing a venue for sustained periodic ecological monitoring at small spatial scales, long duration, and fine taxonomic resolution.

## Introduction

Observational field studies are the foundation of ecology by providing the inspiration and context for experiments, theoretical models, empirical inference, and management. Long-term field studies are disproportionately valuable to ecology and policy (Hughes et al., 2017) and critical to the assessment of biodiversity change in a climate of anthropogenic forcing (Magurran et al., 2010). Most long-term quantitative studies are on the order of several decades (Vellend et al., 2013; Dornelas et al., 2014; Elahi et al., 2015; Kaplanis, 2023), with notable exceptions rooted in the century-long tradition of Western European natural history (Southward, Hawkins & Burrows, 1995; Richardson et al., 2006; Silvertown et al., 2006). Despite their importance, funding for long-term monitoring has declined (Hughes et al., 2017) and thus the maintenance of these studies is a challenge. Besides funding, the transfer of knowledge, methods, and data between sequential investigators is critical to extend long-term datasets beyond a few decades, the typical length of an academic career.

Here we examine the study of a rocky intertidal shore initiated in 1931 and repeatedly sampled thereafter by five different lead investigators or groups (Micheli et al., 2020). In contrast to recent rocky shore monitoring studies (Murray, Ambrose & Dethier, 2006; Kaplanis, 2023), our 90 year historical study trades what are now considered fundamental aspects of study design (e.g., randomization, replication) for the detailed enumeration of the complete macroscopic animal community along a single continuous transect. The 90-year duration is accompanied necessarily by two key investigator-dependent challenges, species identification and variation in sampling effort. We address two primary goals. First, we ask whether temporal change in biodiversity (sensu Magurran, 2021) is sensitive to investigator era. Second, we identify the recent ecological ‘winners’ and ‘losers’, relative to the historical baseline, and ask whether such designations can change over time (1993-2023).

## Methods

### History of the intertidal survey

This study was initiated in 1931 and completed in 1933 by Willis Hewatt, a graduate student at Stanford (Hewatt, 1934). Hewatt surveyed 108 consecutive square yards along a transect perpendicular to shore at Hopkins Marine Station (HMS). A primary focus of Hewatt’s work was the spatial distribution of animals along the shore; the abundances of 90 species in each quadrat were presented in his Table 10 (Hewatt, 1937). Nineteen of these 90 species were not counted but instead categorized qualitatively as “abundant”, “common”, “occasional”, or “rare.” After the surveys had been completed for the 108 quadrats, four “typical squares” were selected as representative of intertidal zones; these four quadrats were resurveyed intermittently during changing seasons. A complete list of species (n = 170) was also provided (Section VII; Hewatt, 1937); Hewatt acknowledged assistance with the identification of crustaceans, gastropods (including nudibranchs), and fishes.

Bruce Provin was the first to resurvey Hewatt’s transect as part of a student project at HMS (Provin, 1949). Provin relocated (using permanent markers and maps) and resurveyed four of Hewatt’s “typical squares” (quadrats 12, 24, 35, 90). All macroscopic organisms were counted and presented in the manuscript; those that couldn’t be counted exactly were rated as “abundant”, “common”, “occasional”, or “rare.” Species were identified using *Between Pacific Tides* (Ricketts & Calvin, 1948), *A Laboratory and Field Textbook of Invertebrate Zoology* (Light, 1941), and various monographs and papers; help was also obtained from the course instructors and other students.

Between 1993 and 1996, Hewatt’s transect was again resurveyed by undergraduate students (Rafe Sagarin, Sara Gilman) and their mentors (Charles Baxter, Jim Barry); hereafter referred to as SBGB (Sagarin et al., 1999). Sagarin and Gilman relocated and resurveyed 57 plots between spring 1993 and summer 1995. During summer 1996 SBGB resurveyed the first 19 plots surveyed in spring 1993; this included quadrats 27-38 and 62-68 (these 19 quadrats are hereafter referred to as ‘core’ quadrats). With few exceptions, all individuals within a plot were counted, including those on or under marine plants or other species. Species that could not be readily and nondestructively identified were not counted; only species that could be identified with the unaided eye were counted (Sagarin et al., 1999). Regional guides were used for species identification (Smith & Carlton, 1975; Morris, Abbott & Haderlie, 1980).

To continue the long-term monitoring effort between 1999 and 2015, Sagarin returned to HMS intermittently to resurvey the core quadrats. These resurveys occurred in 1999, 2002, 2005, 2009, 2014, 2015. No description of methods is available; we assume that the methods were similar to the methods described by SBGB for the years 1993 and 1996. Field notes indicated that within a given year, quadrats were occasionally sampled in different seasons.

After Sagarin’s death in 2015, Fiorenza Micheli and James Watanabe continued surveying the core quadrats on an annual basis. Surveys were conducted between May and July; the sampling effort often included students from an undergraduate spring course at HMS. The sampling methods described by SBGB (Sagarin et al., 1999) were maintained. Regional guides were used for species identification (Morris, Abbott & Haderlie, 1980; Carlton, 2007), with assistance from local experts (e.g., Chuck Baxter, Jim Carlton, John Pearse).

Robin Elahi began leading the survey effort in 2019, with transfer of knowledge from Micheli and Watanabe. In addition to the core quadrats, Elahi began surveying quadrats 12, 16, 20, and 24 (hereafter referred to as ‘extra’ quadrats) to address biodiversity change in the upper intertidal. In addition to the annual late spring sampling, a subset of quadrats was also sampled during winter months. The sampling methods remained the same, with a few exceptions (detailed in Appendix S1). Regional guides were used for species identification (Morris, Abbott & Haderlie, 1980; Carlton, 2007).

We divided the survey efforts into six investigator eras, because we hypothesized that differences in species identification and sampling effort could affect inferences of biodiversity change over the ninety years of sampling. Additional details on sampling methods are provided in supplementary information (Appendix S1), including a visualization of the relative positions, tidal heights, and vertical relief of quadrats along the historical transect (Appendix S1, Fig. S1).

### Harmonizing taxonomy and quantifying taxonomic overlap

The qualitative and quantitative observations for core and extra quadrats across all investigator eras were assembled into a database. For species listed in our database, we tabulated the scientific names used by Hewatt (1937) and Sagarin et al. (1999), reconciling them with current names provided in the World Register of Marine Species (WoRMS Editorial Board, 2023) (Appendix S2, Table S1). For example, the whelk *Acanthinucella punctulata* was recorded as *Acanthina punctulata* and *Acanthina lapilloides* by Sagarin et al. (1999) and Hewatt (1937), respectively.

Once we synonymized the species names in the database, we next tabulated the overlap in species presence across investigator eras (Appendix S2, Table S2). We omitted amphipods (*Ampithoe*, *Atylopsis*, *Melita*) and spirorbid tubeworms (e.g., *Spirorbis*) from this overlap analysis (and subsequent biodiversity analysis) because these species are small, inconspicuous, and the monitoring records suggested inconsistencies in surveying effort for these taxa. We also omitted animals recorded at taxonomic levels above genus, which started in 2020. We then removed animals that were recorded as genera, when there was also a species recorded within that genus. In cases where a species (or genus) was recorded in 1993 or after, but the abundance was not quantified by Hewatt, we checked to see whether Hewatt had recorded the species in the complete list (Appendix X in Hewatt, 1934). If it was not on the list, we recorded it as absent from Hewatt’s study; if it was on the list, then we recorded it as present in Hewatt’s study. In total, 208 animals at the genus or species level were identified across the 90 years of the study (Appendix S2, Table S1). We visualized species overlap for three investigator eras (1931-1933, 1993-1996, and 2020-2023). We then repeated the visualization, but using species for which quantitative abundances were available.

### Inferring biodiversity change across investigator eras

In Table 1, we summarize the necessary steps to prepare the historical database for our primary question – how has biodiversity changed on the historical transect, while considering the different investigator eras? We included years from five (of six) investigator eras when all nineteen core quadrats were sampled, omitting the 1949 era because Provin (1949) sampled only three of these core quadrats. Using the qualitative dataset described above (steps 1 and 2 in Table 1), we then removed hermit crabs (*Pagurus*), and limpet epibionts (*Lottia asmi*, *Lottia* spp.) on trochid snails (*Tegula*) because they were not sampled quantitatively by Hewatt. Next, we had to reconcile potential ambiguities in species identification across investigator eras. For each taxon listed in Table S1 (Appendix S2), we identified the taxonomic level at which we felt comfortable inferring patterns of biodiversity change and summarized the abundances to that level (step 3 in Table S1). Lastly, we removed taxa above genus; all processing steps are summarized in Table S3 (Appendix S2). Total numbers of species, genera, and clades in the quantitative dataset are summarized in Table S4 (Appendix S2).

**Table 1.** Recommended steps for harmonizing taxa across investigator eras for robust inferences on biodiversity change over time.

| Step | This paper |
| --- | --- |
| 1. Synonymize taxa using accepted database (e.g., World Register of Marine Species) to account for changes in taxonomy over time | Table S1 |
| 2. For every taxon, tabulate whether it was observed by each investigator to quantify taxonomic overlap between eras and facilitate the discovery of misidentifications (e.g., within genus) | Figure 1, Table S2 |
| 3. For each taxon, identify the taxonomic level to which it will be grouped for analysis (in our case, genus or species) | Table S1 |
| 4. Identify all the processing steps from the raw dataset to the final dataset used for analysis | Table S3, S4 |
| 5. When testing for temporal trends in alpha or beta diversity, include the effect of investigator era in the statistical model | Table 2 |

Temporal patterns at the quadrat scale (0.84m^2^) were examined for richness (i.e., species density; Gotelli & Colwell, 2001), Hill-Shannon diversity, and Hill-Simpson diversity; these three diversity indices are also known as Hill numbers of orders *q* = 0, 1, and 2, respectively (Hill, 1973). Hill numbers can be interpreted as the effective number of species or genera (i.e., unique taxa) for all taxa (*q* = 0; richness), common taxa (*q* = 1; Hill-Shannon), and dominant taxa (*q* = 2; Hill-Simpson) (Chao et al., 2014). The use of these three metrics (rather than just one) permits a more complete story about biodiversity change that incorporates nuances related to rare versus abundant species, and their relative importance.

For the quadrat scale, diversity indices were averaged across all quadrats (n = 19) and visualized with 95% confidence intervals (assuming a normal distribution). Although the same 19 quadrats were sampled in each of the 18 years, the number of individuals within quadrats was highly variable and could confound interpretations of changes in diversity per se (Gotelli & Colwell, 2001). Therefore, we also evaluated temporal patterns in diversity at the site level using a coverage-based estimator, which better accounts for the underlying species abundance distribution of the community being sampled than traditional rarefaction (Hsieh, Ma & Chao, 2016; Roswell, Dushoff & Winfree, 2021). Hereafter, we will refer to the number of taxa (at quadrat and site scales) as ‘richness’, and Hill-Shannon and Hill-Simpson indices as ‘diversity’.

In addition to examining temporal variation in alpha diversity metrics, we investigated temporal trends in beta diversity using the same quantitative database that was used for richness and diversity. We tested the hypothesis that biotic homogenization (i.e., increasing spatial similarity) has occurred on the transect over the past three to nine decades, by quantifying temporal patterns in spatial beta diversity (Olden & Rooney, 2006). For each year, we calculated the median pairwise Jaccard distance (based on species presence-absence) between each quadrat and every other quadrat and then averaged these quadrat medians to obtain an estimate of similarity at the site scale; in this framework a positive temporal trend in community similarity indicates biotic homogenization.

We used Spearman rank tests to assess the significance of temporal correlations of alpha diversity metrics and spatial beta diversity over the entire course of the study (1933-2023). We used rank correlation tests because the historical data, collected by Hewatt in 1931-1933, preceded the remainder of the data by 60 years and thus could potentially exert considerable leverage in a linear regression. The Spearman rank tests necessarily omitted the effect of investigator era. However, we used linear models (analysis of variance; ANOVA) to test the effects of time (year) and investigator era (era) of alpha diversity and beta diversity over the final 30 years of the study (1993-2023). We did not incorporate temporal autocorrelation into our linear models because the residuals did not exhibit evidence of a time lag for any of the diversity metrics. Residuals of the linear models did not display patterns of heteroscedasticity.

### Identifying ‘winners’ and ‘losers’ over time

In studies of biodiversity change, species that are more or less abundant than they used to be sometimes are defined as ‘winners’ or ‘losers’, respectively. We used the quantitative database to first identify winners and losers. Second, we asked whether the designation of winners and losers varied over time, by leveraging the time-series data from 1993-2023. To summarize changes in the composition of animal taxa on the historical transect, we compared each year of sampling in the recent thirty years (1993-2023) to the historical baseline (1931-1933). First, the total abundance of each genus (across 19 quadrats) was calculated for each year. Then the log change in abundance was calculated for each of the recent years, relative to the historical baseline (*log change* = *ln*[(*n_year_* + 1) / (*n_baseline_* + 1)]). The average log change and 95% confidence interval (CI) summarized the recent years relative to the historical baseline and identified winners and losers. These designations were examined for temporal consistency by visualizing the trend in log change over the recent thirty years.

### Software and data

We relied on the ‘tidyverse’ (Wickham et al., 2019), ‘iNext’ (Hsieh, Ma & Chao, 2016), ‘vegan’ (Oksanen et al., 2011), ‘eulerr’ (Larsson et al., 2016), and ‘patchwork’ (Pedersen, 2024) packages in R 4.3 (R Development Core Team, 2023) for data processing, visualization, and analysis. All data and code are available in a permanent repository (https://purl.stanford.edu/fd733kb4905).

## Results

### Harmonizing taxonomy and quantifying taxonomic overlap

We checked our species names with those used by Hewatt (1937) and Sagarin and colleagues (Barry et al., 1995; Sagarin et al., 1999). After harmonizing the taxonomy to the presently accepted names (Appendix S2, Table S1), the database contained 232 unique taxa (180 species, 28 genera, 24 higher level clades), of which 121 species were observed in more than one era and thus could have been subject to a name change. Of these 121 species, 42 species changed names once (35%), and 12 species changed names twice (10%).

The taxa recorded on the historical transect varied considerably between each of the six investigator eras (Appendix S2, Table S2). For example, only 24% of the taxa (genera and species) was shared by three eras (1931-1933, 1993-1996, 2020-2023) for the qualitative (i.e., complete) set of animal taxa (Fig. 1A), and the quantitative subset of animal taxa (Fig. 1B). The most recent era (2020-2023) shared 36-44% of taxa with 1993-1996, and 29-32% of taxa with 1931-1933. Taxonomic uniqueness within these three eras ranged from 6 to 32% for the quantitative subset of data. Much of this variability was a consequence of variation in repeated annual sampling and total counts between the eras (Appendix S2, Table S5).

**Figure 1.**
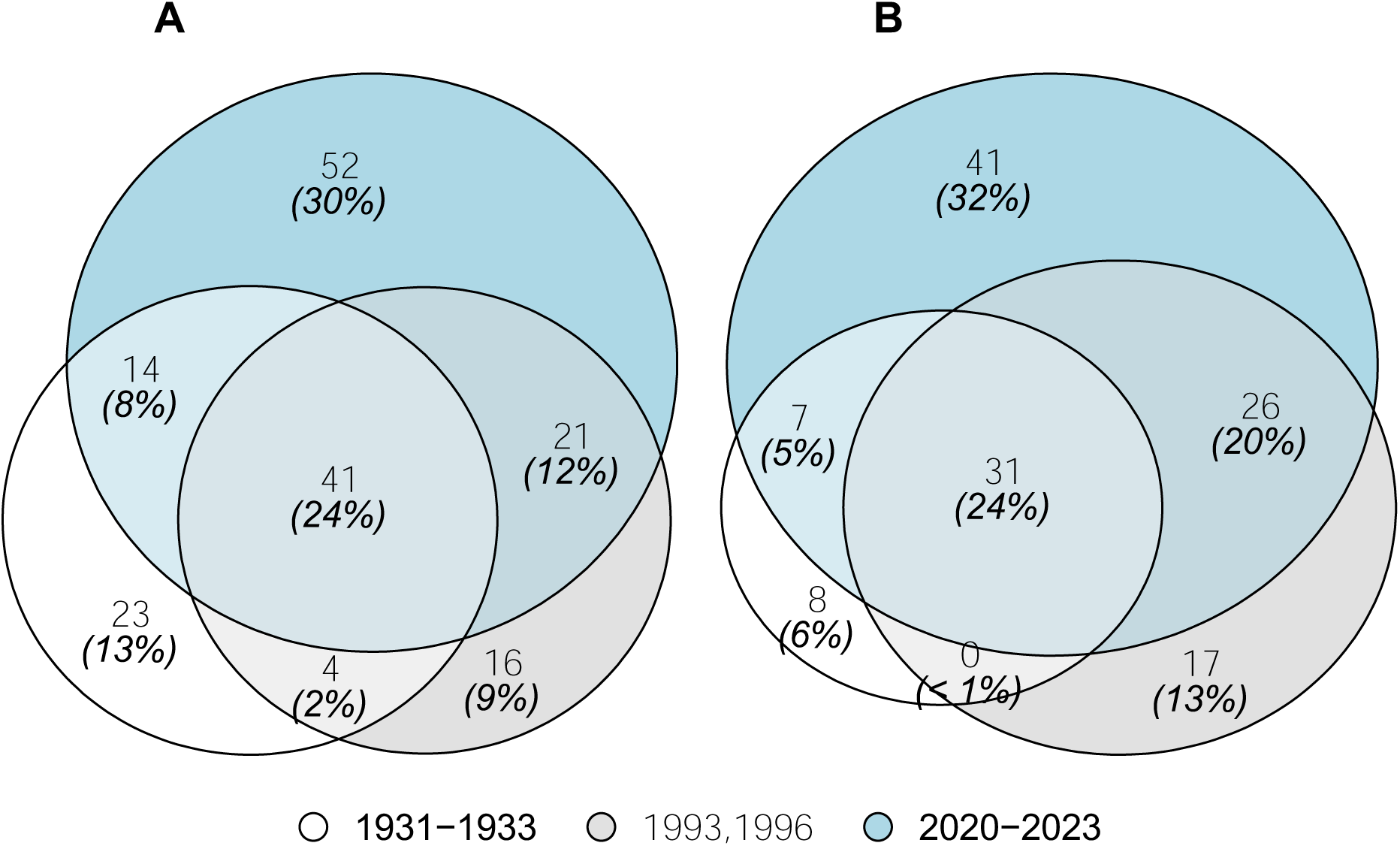
(A) Euler diagrams showing the overlap in the complete set of animal taxa (n = 171) and (B) the quantitative subset of taxa (n = 130) observed on the historical transect during three investigator eras.

In all, 365,914 invertebrates were counted and identified. After removing amphipods, *Spirorbis*, limpet epibionts on *Tegula* spp., *Pagurus* spp., and taxa above genus, 347,543 records remained (95%). After grouping species to higher taxonomic levels for the purposes of quantitative analysis, the wrangled database contained 157 taxa (119 species, 38 genera); this database was used for all temporal analyses of biodiversity change using alpha and beta diversity indices. The wrangled database contained 144 unique animal genera, and a subset of these genera were used to visualize changes underlying the aggregate metrics of alpha and beta diversity. That is, we focused on the genera that comprised 99% of the individuals counted on the historical transect over the nine decades; this threshold removed 108 genera that made up <1% of the individuals, leaving 36 genera.

### Inferring biodiversity change across investigator eras

First, we asked whether 90-year trends in diversity (Figs. 2, 3) were apparent – without considering investigator eras – using Spearman rank tests (Appendix S2, Table S6). At the quadrat scale, Hill-Shannon diversity was correlated negatively with year (ρ = -0.71, *p* = 0.002), and Hill-Simpson diversity was correlated negatively with year (ρ = -0.77, *p* < 0.001). At the site scale, estimated Hill-Shannon diversity was correlated negatively with year (ρ = -0.50, *p* = 0.04); Hill-Simpson diversity was correlated weakly with year (ρ = -0.42, *p* = 0. 08). Richness did not exhibit a temporal trend over 90 years at either spatial scale (Appendix S2, Table S6). Quadrats became more similar to one another, because spatial similarity (i.e., beta diversity) was correlated positively with year (ρ = 0.82, *p* < 0.001).

**Figure 2.**
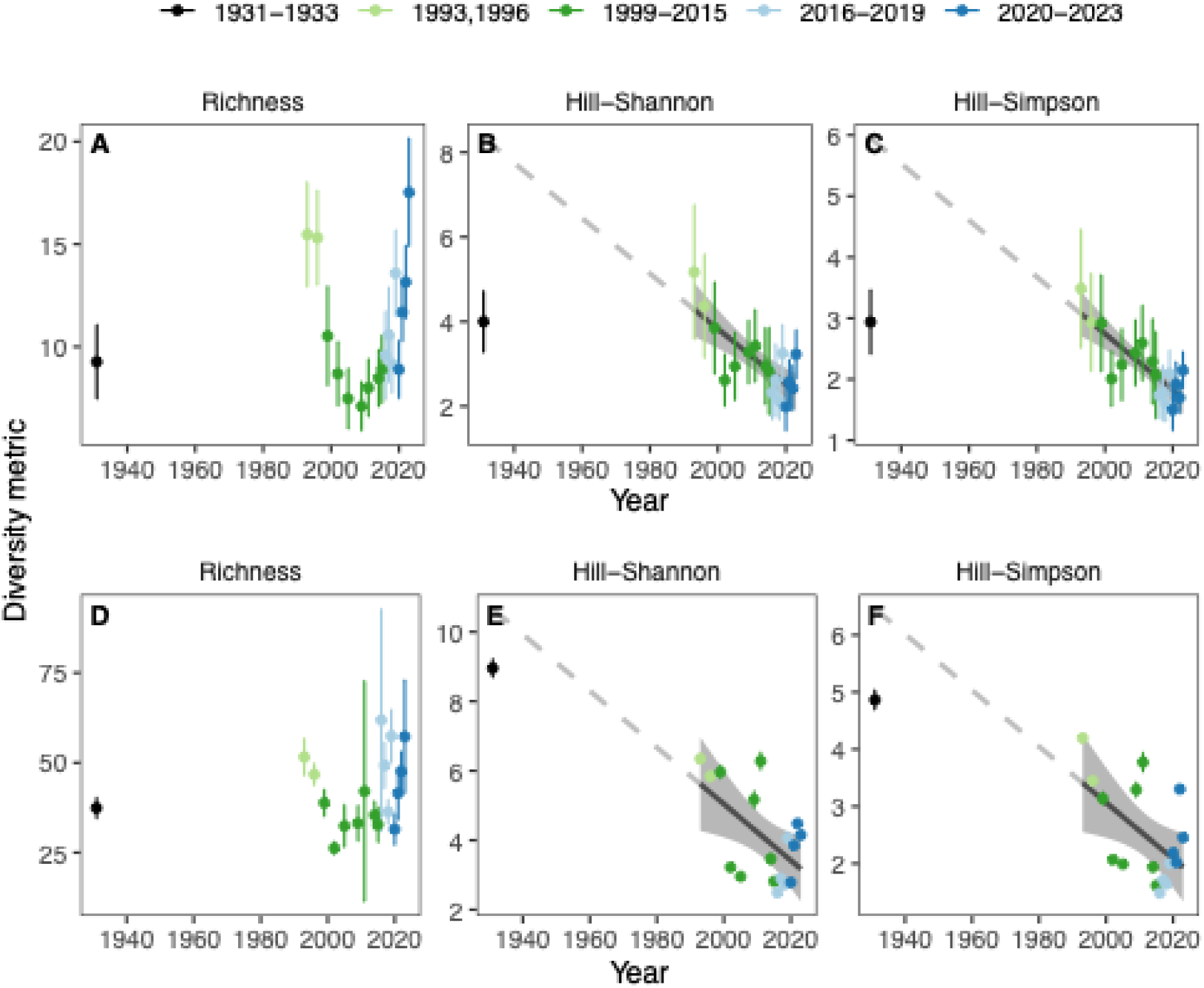
The animal community on the historical transect has declined linearly in diversity over time, but no trends in species richness were observed. (A) Species richness, (B) Hill-Shannon diversity, and (C) Hill-Simpson diversity at the quadrat scale were summarized across quadrats (n = 19; mean ± 95% CI). (D) Species richness, (E) Hill-Shannon diversity, and (F) Hill-Simpson diversity at the site scale were estimated using a coverage-based estimator (estimate ± 95% CI). Linear fits with 95% confidence intervals are visualized for 1993-2023 when the temporal effect was significant (using ANOVA; Table 2). The dashed gray line extrapolates the linear trend to 1933.

**Table 2.**
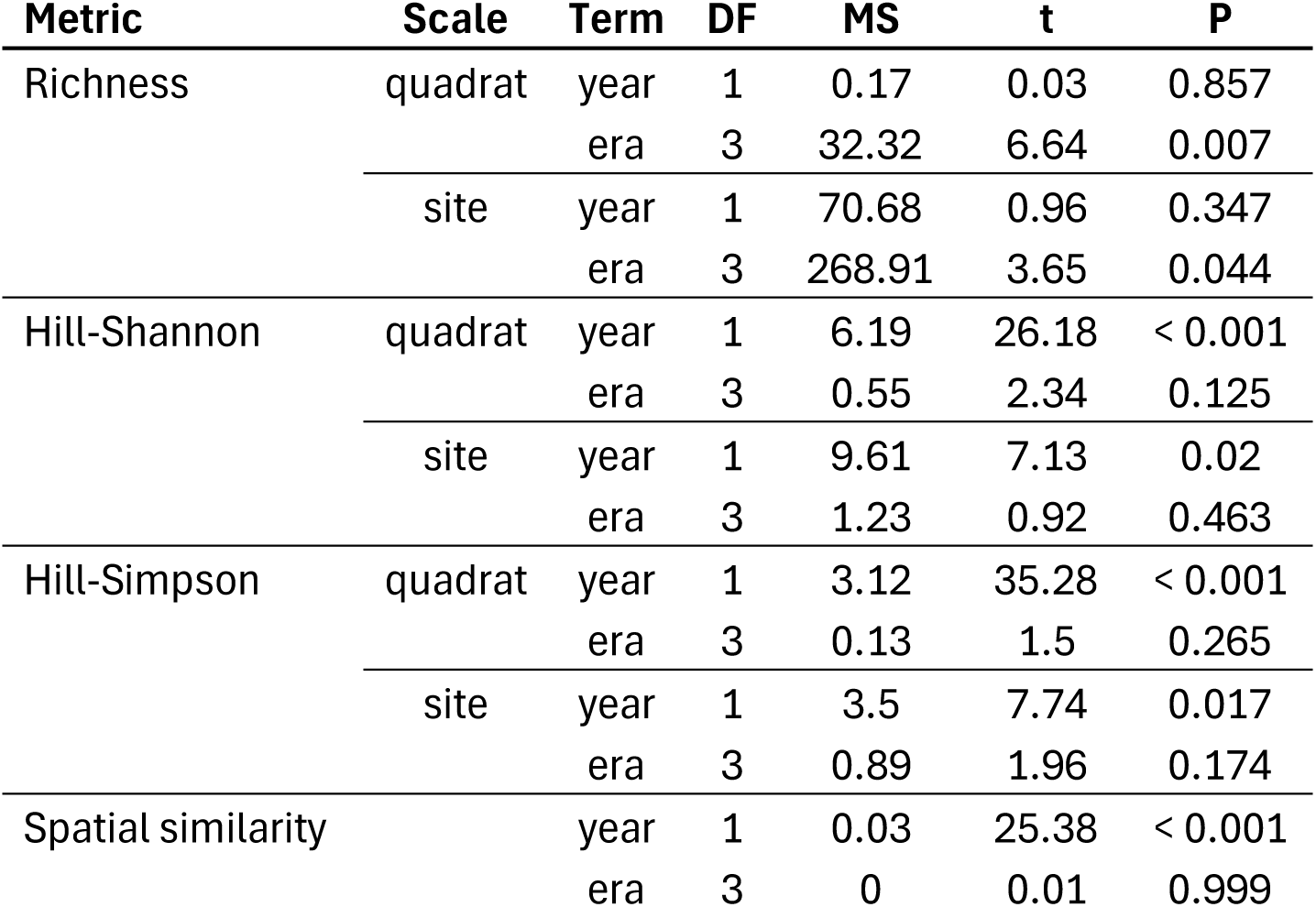
Results of analyses of variance testing the effects of year and investigator era on the biodiversity of the animal community (1993-2023). Biodiversity metrics included alpha (richness, Hill-Shannon, Hill-Simpson) diversity at quadrat and site scales, as well as beta diversity (spatial similarity).

Second, we asked whether linear trends in biodiversity over the recent 30 years could be inferred in the context of investigator eras using ANOVA. For richness, temporal trends were not significant, but the effect of era was significant at both spatial scales (Table 2). For example, richness exhibited a consistent decline between 1999 and 2015 (Fig. 2A, 2D). During this era, species-accumulation curves saturated at relatively low values of site-scale richness (< 50 taxa; Appendix S2, Fig. S2) because fewer individuals were counted, particularly for mobile species (Fig. S3). Counts of sessile taxa exhibited boom and bust dynamics driven primarily by high barnacle densities (which were necessarily subsampled and then extrapolated to the entire quadrat).

In contrast, both Hill diversity indices exhibited linear declines over time at both quadrat and site scales; the effect of era was not significant (Table 2, Fig. 2). At quadrat scales, diversity was comparable between 1931-1933 and 1993-1996 (Fig. 2B, C), but at site scales diversity was higher in 1931-1933 relative to 1993-2023 (Fig. 2E, F). These trends in alpha biodiversity (Fig. 2) represented both sessile and mobile taxa. When split by motility (Fig. S4), mobile taxa exhibited similar temporal trends to Figure 2 (all taxa) but sessile taxa demonstrated no temporal trends, indicating that most of the observed variation over time is due to mobile taxa.

For spatial similarity (i.e., spatial beta diversity), there was no effect of investigator era (Table 2). However, there was a strong linear increase in spatial similarity (Fig. 3) indicating taxonomic homogenization among the quadrats within the historical transect.

**Figure 3.**
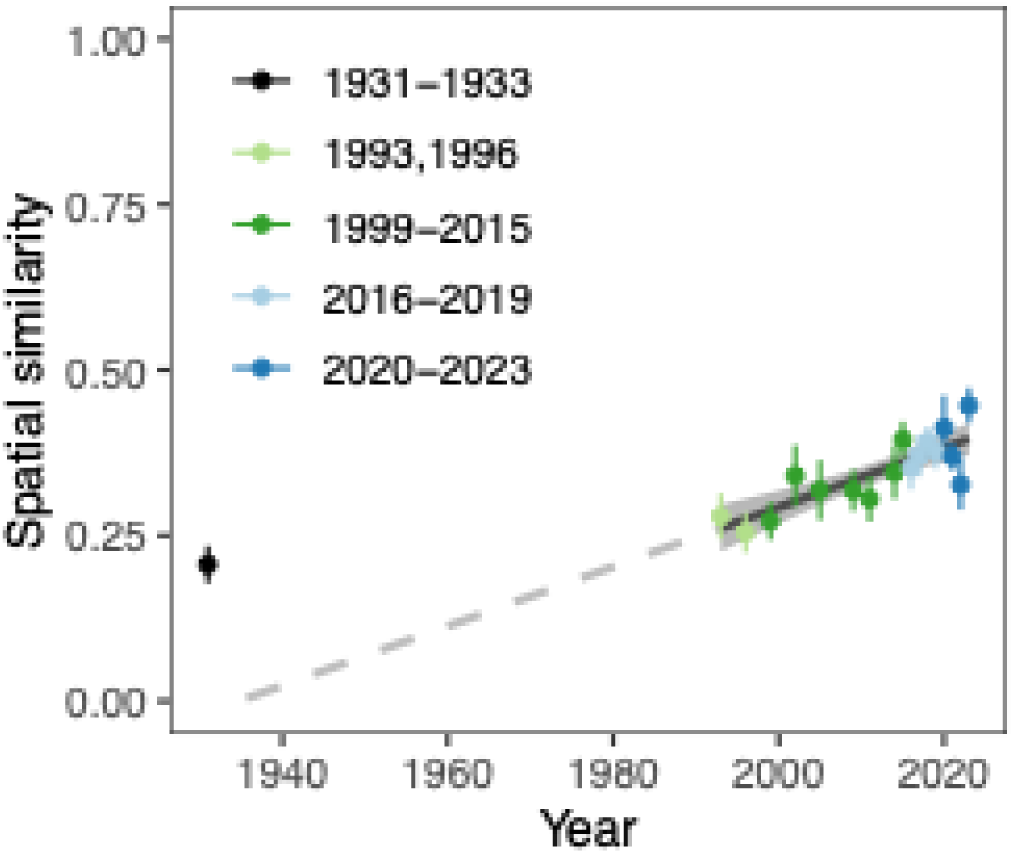
The animal community on the historical transect has become more homogeneous over the past three and nine decades. Similarity was measured using the Jaccard index, where 0 and 1 represent completely different and exactly identical communities, respectively. Linear fits with 95% confidence intervals are visualized for 1993-2023. The dashed gray line extrapolates the linear trend to 1933.

### Identifying ‘winners’ and ‘losers’ over time

We focused on 36 genera that comprised 99% of the total counts for the analysis of recent winners and losers, relative to the historical baseline. The average density of each genus, by era, is provided in Table S7. A significant change in abundance, relative to the original survey by Hewatt in 1931-1933, was observed in 35 of 36 genera at the 95% level, and 33 genera at the 99% level (Fig. 4). Changes in the snail *Littorina* were not significant, and changes in the ascidian *Eudistoma* and isopod *Cirolana* were significant at the 95% level. Of 10 sessile genera, eight were winners; barnacles (*Chthamalus, Balanus, Tetraclita*), tubesnails (*Thylacodes*), anemones (*Anthopleura, Corynactis*), and compound ascidians (*Aplidium*, *Eudistoma*) have become more abundant, while mussels (*Mytilus*) and the social ascidian *Clavelina* have become less abundant over the past 30 years. Mobile invertebrates exhibited a variety of trends. For example, limpets (*Lottia*, *Acmaea*, *Fissurella*) and columbellid gastropods (*Alia, Amphissa*) have declined in abundance, whereas whelks (*Paciocinebrina*, *Acanthinucella*) have increased in abundance. Crabs (*Pachycheles, Pachygrapsus, Petrolisthes, Pugettia*), annelid worms (*Halosydna*, *Phascolosoma*), and echinoderms (*Amphipholis, Leptasterias, Strongylocentrotus*) have declined since the 1930s. Despite these average changes in abundance, many of these taxa exhibited non-linear trends over the recent thirty years. For example, mussels (*Mytilus*) and urchins (*Strongylocentrotus*) have recently recovered to or exceeded historical abundances. In contrast, the strawberry anemone (*Corynactis*) declined abruptly to zero in the last decade (Fig. 5).

**Figure 4.**
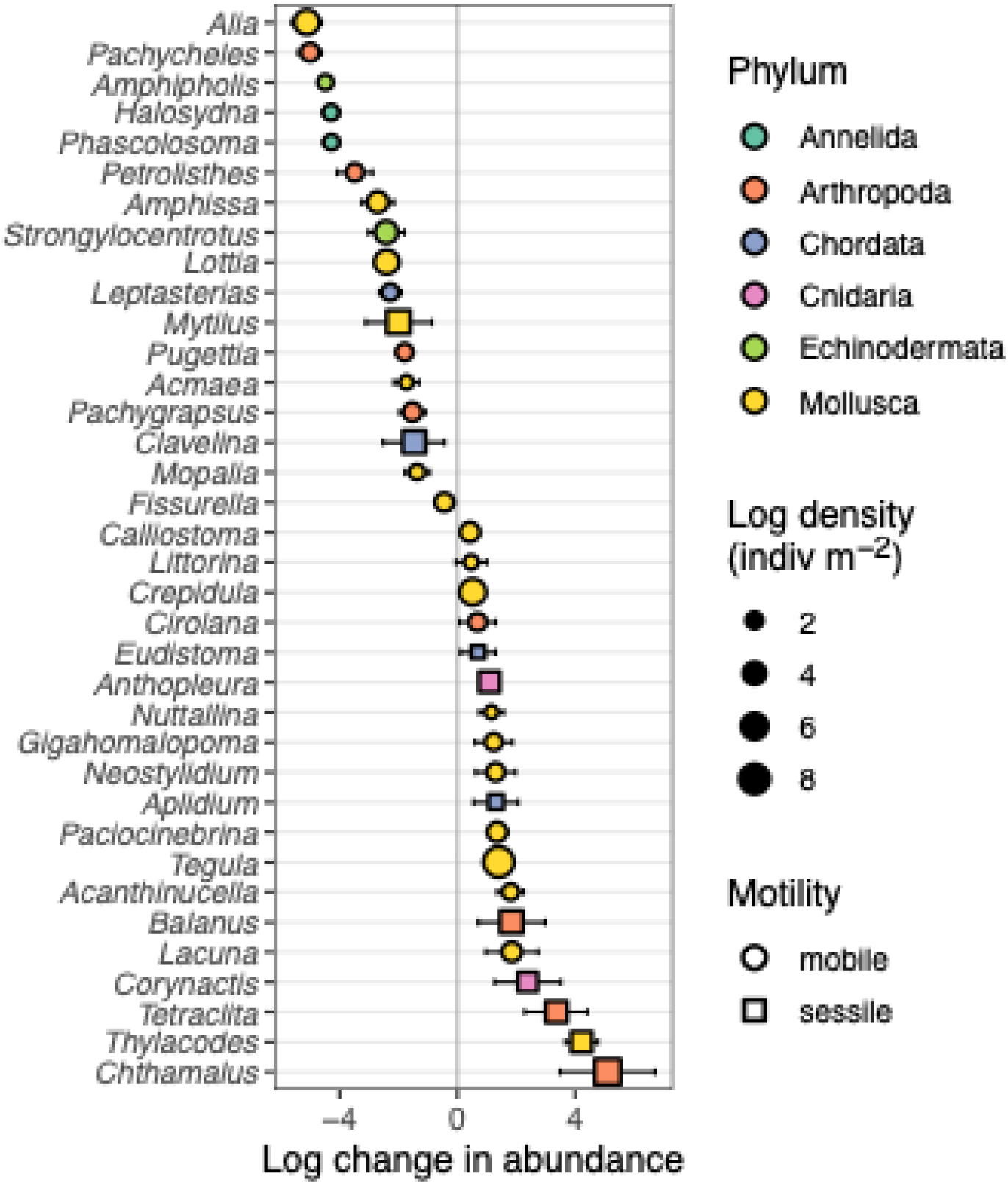
Changes in abundance (mean ± 95% CI; n = 17) of animal genera on the historical transect over the past 30 years (1993-2023) relative to the initial survey in 1931-1933. Change in abundance (the total abundance for each genus in year *i*, where *i* ranges from 1993 to 2023) is expressed as the natural log of a ratio [ln(abundance + 1 in year *i* / abundance + 1 in the original survey). The size of points corresponds to the log mean density, averaged across all 18 years of sampling (1931-1933, 1993-2023). These 36 genera comprise 99% of the individuals counted on the transect over the 90-year duration of the study.

**Figure 5.**
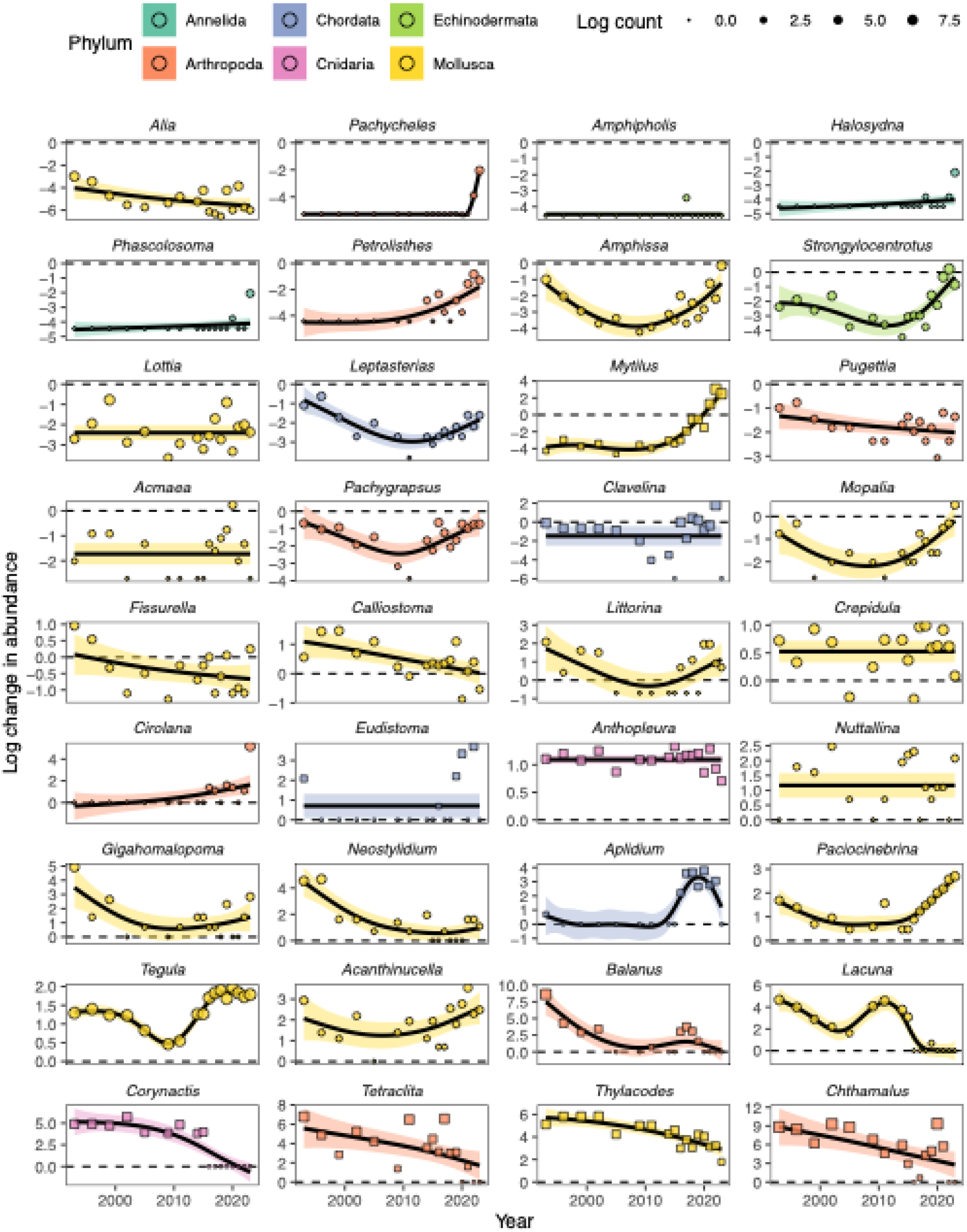
Temporal variation (1993-2023) in the log change in abundance of animal genera on the historical transect, relative to the historical baseline in 1931-1933. Change in abundance (the total abundance for each genus in year *i*, where *i* ranges from 2020 to 2023) is expressed as the natural log of a ratio [ln(abundance + 1 in year *i* / abundance + 1 in the baseline year)] where 1931-1933 was the first sampling event. The size of points corresponds to the log total count. These 36 genera comprise 99% of the individuals counted on the transect over the 90-year duration of the study. Squares and circles represent sessile and mobile genera, respectively. Smoothed curves represent generalized additive models.

## Discussion

Long-term monitoring studies must inevitably ‘pass the quadrat’ from one investigator to the next. Using an intertidal case study spanning three decades and six lead investigators, we first highlight the aspects of biodiversity change susceptible and robust to investigator effects. Second, we emphasize that inferences about ecological ‘winners’ and ‘losers’ can be complicated by non-linear population trajectories especially when time series are truncated. We conclude with recommendations based on the lessons we have learned from field sampling and the downstream analysis of the data.

One of the strengths of our intertidal sampling program, initiated by Hewatt (Hewatt, 1937) and restarted in earnest by Sagarin and Gilman (Barry et al., 1995; Sagarin et al., 1999), is the enumeration of all macroscopic invertebrates in our quadrats. This feature permits the estimation of biodiversity metrics (e.g., richness, diversity), which rely on (i) quantifying abundance, and (ii) maintaining on ‘open’ species list, that is, a list that permits the addition of new species when discovered. Given the high density of intertidal invertebrates on the rocky shore, and the taxonomic expertise required to identify taxa, it is imperative to examine whether inferences on biodiversity change are sensitive to observer effects. In the context of sampling small and numerous rocky shore invertebrates, observers may vary in their taxonomic ability, eyesight, time constraints, attention to detail, and access to species identification references; these aspects can also change over time for the same observer. During each sampling event in our case study, the lead investigator may enlist the help of volunteers and students to assist with the identification and counting. Ultimately, it is the responsibility of each lead investigator to ensure quality control with respect to sampling and species identification, and thus we have focused on six distinct investigator eras. We acknowledge that the actual observers add another layer of uncertainty to biodiversity estimates, but that is beyond the scope of the present study.

Our analyses suggest that investigator effects are particularly relevant for species density (number of species observed in a quadrat or sample) and species richness (number of species estimated at the site scale from quadrats). These metrics hinge on the relative abundance of species and the identification of rare, cryptic and/or potentially new species. Species density is well known to be subject to biases related to sampling effort (Gotelli & Colwell, 2001), and we observed trends in species density to be strongly non-linear and associated with different investigator eras. Estimated species richness (at the site scale), instead displayed no trend over time but was still influenced by investigator era. The lack of a linear temporal trend in our case study is consistent with theory suggesting a net balance between species colonizations and extinctions over time (Brown et al., 2001) and recent syntheses that emphasize high variability in local-scale biodiversity trends (Vellend et al., 2013; Dornelas et al., 2014; Elahi et al., 2015).

In contrast to richness, diversity metrics (i.e., Hill-Shannon and Hill-Simpson) that reduce the leverage of rare species (Roswell, Dushoff & Winfree, 2021) were robust to investigator effects in our study. Across quadrat and site spatial scales, the diversity of common (Hill-Shannon) and dominant (Hill-Simpson) species has declined over the past three decades and to a lesser extent since the 1930s. Temporal declines in Hill indices were accompanied by increases in spatial similarity. That is, the quadrats across the transect became more similar to each other over time, indicating biotic homogenization (Olden & Rooney, 2006; Blowes et al., 2024). Notably, these biodiversity trends were driven primarily by mobile invertebrates, rather than sessile invertebrates. The latter were less speciose but exhibited large inter-annual population fluctuations, thus contributing to temporal variability but no directional trend in diversity indices. In contrast, mobile taxa exhibited strong linear declines in alpha diversity. We found it surprising that mobile taxa exhibited a stronger signal of biodiversity change over time, considering that they can move in and out of quadrats and thus respond to the environmental conditions on any given sampling day. However, this potential artifact may indeed be smaller than the variation induced by the extreme (and visually apparent) recruitment potential of sessile barnacles and mussels. Moreover, many of the sessile species were clonal and the method of counting individuals does not reflect the ecological process of occupying space (e.g., for sponges or compound tunicates). To complicate matters, individual zooids/polyps were counted for *Clavelina* and *Corynactis*, but in fact these counted units do not represent genetically distinct individuals. Photographs of the quadrats (taken since 2015; Elahi & Watanabe, 2023) can instead be used to assess future dynamics of the sessile community using percent cover, rather than counts, as an ecologically relevant estimate of abundance.

Our biodiversity analyses demonstrate that the animal communities on the historical transect have become simpler and more similar, but rare species can still be found if enough effort is expended by the observer. These quantitative results align with recent oral histories by researchers and professors that taught or lived on the Monterey Peninsula for more than five decades (Pearse et al., 2017). In particular, anecdotes from Charles Baxter (Lecturer, Hopkins Marine Station), John Pearse (Professor, UC Santa Cruz), and Steve Webster (Senior Marine Biologist, Monterey Bay Aquarium) suggest that at the start of their careers as marine biologists (in the 1960s) the diversity of life in the intertidal was “absolutely incredibly abundant diverse and everything” (C. Baxter); “you would turn over a rock … and there were uncountable numbers of brittle stars, isopods, porcelain crabs” (S. Webster); “the diversity there and the number of species and the colors, I never got over it. I was always coming back to Pacific Grove to look at the diversity” (J. Pearse). However, towards the end of their careers, it became harder to find certain species (e.g, for teaching invertebrate zoology courses), and in general, the color and diversity of intertidal life was muted: “in the 80s I began to notice the intertidal looked different … it was a change that began to make it look more like it did in Southern California than it did up here.” (C. Baxter); “you just don’t see those [cucumbers] anymore. Or if you see them, much, much smaller numbers of them. My last trip to the intertidal I think I saw three brittle stars. Well in the old days you’d turn one rock over, you’d see 30 of them.” (S. Webster); “we used to see lots and lots of polyclad flatworms … it’s not like they’re gone, you just have to spend time looking. There used to be … a little carnivorous isopod, *Cirolana*. It used to be that, I remember, you turn over the rock and they would just scurry, just explode. It’s nothing like that now.” (J. Pearse). The alignment of expert memories from the Monterey Peninsula with our quantitative analyses corroborates the general conclusions regarding diversity decline in the intertidal.

There were some clear ‘winners’ and ‘losers’ underlying changes in alpha and beta diversity on the historical transect. Here we define winners and losers as those genera with consistently higher and lower densities, respectively, in the recent 30 years relative to the 1930s. Winners included certain snails (*Tegula*), anemones (*Anthopleura*), chitons (*Nuttalina*), barnacles (e.g., *Chthamalus*, *Tetraclita*), and whelks (e.g., *Acanthinucella*, *Paciocinebrina*). Losers included other snails (*Alia*), limpets (*Lottia*, *Acmaea*), crabs (*Pugettia, Pachygrapsus*), and brittle stars (*Amphipholis*). However, many of the apparent winners and losers displayed non-linear trajectories over the past thirty years. In particular, mussels (*Mytilus*) and urchins (*Strongylocentrotus*) recovered to or exceeded historic levels after prolonged declines; urchin recruits were particularly abundant in the dense mussel beds towards the wave exposed portion of the transect after 2020. An increase in the deposition of mussel shells and sand, associated with intense winter storms in 2022 and 2023 may have facilitated the recovery of cryptic species (*Pachycheles*, *Phascolosoma*, *Petrolisthes*, *Cirolana*, *Halosydna* and other polychaete worms) that occupy interstitial and mussel shell habitats during those years. Distinguishing between a population rebound associated with an increase in favorable habitat versus investigator effort remains a challenge for these cryptic and less common species. More generally, the assignment of labels such as winners and losers based on two time points is problematic in the context of dynamic population trajectories, and again highlights the value of long-term periodic monitoring efforts (Hughes et al., 2017).

Despite the prevalence of nonlinear trends, there were several unambiguous changes in the community relative to the 1930s. First, the intertidal green anemones with southern affinity (*Anthopleura sola*) have become well-established on the transect. Second, the smaller *Chthamalus* and southern volcano barnacle (*Tetraclita rubescens*) have replaced the formerly more abundant *Balanus glandula*. Associated with these barnacle increases, whelks have increased (*Acanthinucella punctulata* is another species with southern affinity). Lastly, *Tegula* snails have replaced *Lottia* limpets as the dominant grazer along the transect.

It is beyond the scope of our study to ascribe mechanisms to the many changes in biodiversity and species composition, but we hypothesize that the conspicuous patterns are consistent with predictions related to climate warming. That is, we expected to see increases in the abundance of species with southern affinity (Sagarin et al., 1999; Sanford et al., 2019; Micheli et al., 2020) and increases in the prevalence of smaller-bodied species and individuals (Forster, Hirst & Atkinson, 2012; Ohlberger, 2013; Elahi, Miller & Litvin, 2020). We speculate that the shift from limpet to snail grazers is related to their morphology. Given their large foot and inability to retract within their shell, the body temperatures of limpets are more likely to match surrounding rock temperature which may influence survival during periods of thermal stress (Denny & Harley, 2006; Miller & Denny, 2011). The reduction in limpets and other mobile invertebrates may also be related to the loss of large brown rockweeds (*Silvetia compressa*) on the historical transect (Sagarin et al., 1999); these seaweeds provide a large canopy that ameliorates thermal and desiccation stress. This hypothesis highlights a weakness of the historical program, which does not presently quantify macroalgae in the field. However, we began photographing the quadrats in 2015 (Elahi & Watanabe, 2023) and this will allow us to test hypotheses about the sessile invertebrate winners, and their algal competitors, in the future. Despite the above hypothesized long-term consequences of warming, we did not observe a short-term effect of the 2014-2016 marine heat wave. This appears to be consistent with a broader pattern of recent stability of intertidal rocky shores in central California (Miner et al., 2021), as well as northern California and Oregon (Gravem et al., 2024).

A major limitation of our long-term study is that the animals were sampled from contiguous quadrats along a single transect that may not be representative of other rocky shores. We view this as a worthwhile tradeoff with the duration of the study and the level of taxonomic detail. Such trade-offs in design and implementation are common in other long-term rocky shore studies (Kaplanis, 2023). As a stark contrast to our study, a spatially extensive, well-replicated, and coordinated effort across the west coast of North America must, out of necessity, limit surveys to a pre-selected group of focal species; more thorough taxonomic surveys can only be completed on a less regular basis when funding permits (Multi-Agency Rocky Intertidal Network; Gilbane et al., 2022). Our historical resurvey with its open species list highlights the value of marine stations that enable otherwise unfundable monitoring efforts at small spatial scales but fine taxonomic resolution and long duration.

Nevertheless, we see room for improvement in our approach to the historical study that does not necessarily require more effort, which would be untenable without funding. Instead, a reallocation of current effort would permit stronger inferences about the patterns and underlying causes of future biodiversity change. For example, by focusing on a subset of the existing quadrats, and selecting new permanent quadrats stratified along an intertidal gradient, we could achieve randomization and representation of the local rocky shore. Within the permanent quadrats, a priority is installing temperature loggers to begin relating variation in thermal microclimates to biodiversity changes over time. Increasing the numbers of photoquadrats on the rocky shore would allow the documentation of changes to functional groups of sessile invertebrates and seaweeds with stronger replication but minimal field effort. Many of the trade-offs in sampling design are well known (Murray, Ambrose & Dethier, 2006), but implementing new strategies in the midst of a historical survey will require care.

In summary, there are several key recommendations from our intertidal sampling program that are relevant to inferences about biodiversity change from long-term ecological monitoring studies. First, it is critical to have an open species list, and an open mind to recognize unusual animals that may be new colonizers, especially in the context of climate change and species range shifts (Sagarin et al., 1999). Identifying species to the lowest level possible is difficult but valuable, even if a particular analysis necessitates the grouping of species within genera. Second, consistency in the number of samples (e.g., quadrats) does not guarantee consistency in species abundances, which we interpret here as variation in sampling effort by investigators. Therefore, inferences about trends in species richness can be sensitive to investigator effects because of the inherent difficulty in observing rare or cryptic species. In contrast, diversity indices that deemphasize rare species may be more reliable indicators of biodiversity change in long-term studies that change hands. Third, integrating sampling efforts with the teaching of field courses is one way to maintain a small-scale monitoring effort over time. Moreover, collaborative field courses have a disproportionate impact on students’ sense of belonging to science in general and ecology in particular (Race, Beltran & Zavaleta, 2021). Feedback from our recent students highlights the value of being in the field and contributing to a ninety-year study to understand biodiversity change on a rocky shore.

## Acknowledgements

We are grateful to the previous investigators, as well as many students and volunteers, who contributed their time in the field to revisit and maintain the long-term monitoring effort on the historical transect at Hopkins Marine Station. We thank G. Donahue for feedback on an early version of the manuscript.

## Notes

https://purl.stanford.edu/fd733kb4905

### Competing Interest Statement

The authors have declared no competing interest.

