## Appendix 1 for "Passing the quadrat: inferring biodiversity change over time and across investigators"

**Appendix S1.**

**Section S1. Supporting Methods: description of survey methods for the Hewatt transect, Hopkins Marine Station, California, USA**

Description of methods, 1931-1933 (Hewatt)

The following descriptions of the original survey methods were paraphrased from Willis G. Hewatt's PhD dissertation (Hewatt, 1934) and resulting manuscript in the American Midland Naturalist (Hewatt, 1937). No personal interpretations were included unless explicitly noted.

The study site is a rocky (granite) intertidal shore at Hopkins Marine Station, Cabrillo Point, California, USA. In 1930 Professor G.E. MacGinitie established four benchmarks at intervals along a line perpendicular to shore, across the entire intertidal area directly north of the Jacques Loeb Laboratory. These benchmarks were established at 5.60, 1.16, 0.51, and 0.09m above mean lower low water (MLLW). These benchmarks were installed in collaboration with the U.S. Coast and Geodetic Survey.

In October 1931, W.G. Hewatt initiated a quantitative and qualitative survey of 108 square yards along the permanent transect. A wooden frame (1 yard$^2$; 0.84m$^2$), was placed serially along the west side of the line. A topographic sketch was made of each square quadrat, and the number of individuals of each species was recorded for each quadrat. A stadia rod and leveling instrument was used to sketch the vertical contours of the transect. The goal of Hewatt's research was to "determine more definitely the nature and causes of the vertical and horizontal zonation of the littoral invertebrates of the rocky shore".

A primary focus of the research was concerned with the spatial distribution of 90 species along the transect (Table 10 in Hewatt, 1937). Nineteen of these 90 species were not counted but instead categorized qualitatively as "abundant", "common", "occasional", or "rare." After the surveys had been completed for the 108 quadrats, four quadrats were selected as representative of intertidal zones; these four quadrats were resurveyed intermittently during changing seasons. The work was completed in June 1933, and additional observations were made during the summer of 1934. A complete list of species (n=170) was also provided (Section VII in Hewatt, 1937); Hewatt acknowledged assistance with the identification of crustaceans, gastropods (including nudibranchs), and fishes.

Description of methods, 1947 (Provin)

The following descriptions of the first resurvey of Hewatt's transect were paraphrased from B. Provin's research paper from a Stanford class (Provin, 1949). No personal interpretations were included unless explicitly noted.

Proving resurveyed four of Hewatt's "typical squares" (12, 24, 35, 90); these were described by Hewatt as representative of four intertidal zones. Provin was able to relocate the first three benchmarks; the fourth in the mussel bed was not relocated but was unnecessary to "definitely locate the successive square yard areas to within an error of certainly no more than six inches and probably less than this." All macroscopic organisms were counted; those that couldn't be counted exactly were rated as "abundant", "common", "occasional", or "rare." Species were identified using *Between Pacific Tides* (Ricketts & Calvin, 1948), *A Laboratory and Field*

*Textbook of Invertebrate Zoology* (Light, 1941), and various monographs and papers; help was also obtained from instructors and other students.

For quadrat 12, all collecting and counting was done in one day (about four hours). In this quadrat, the sand and shell fragments were combed through to a depth of about a half inch. For quadrat 24, all collecting and counting in the field was done over four days (about 12 hours total). For quadrat 35, all collecting and counts in the field were completed over two days (about 8 hours). All individuals of the majority of species were taken in the laboratory for identification and counting. Shell and sand was combed through to a depth of a half inch; a rock was turned over and organisms on its underside were also counted. In this square, it was noted that "other annelids – of several kinds – were common in the sand" – but not enumerated. For quadrat 90, all collecting and counting in the field occurred over two days (about 5-6 hours total). The top two to three inches of sand and shell was "dredged" and put into a bucket. All animals and plants were removed from the rocks and crevices and later identified in the laboratory (which took three full days and two evenings). This involved combing through and pulling apart seagrass roots, algal holdfasts, and invertebrate colonies to search for macroscopic organisms. In this square, it was noted that "several syllids were found and annelids of other kinds were abundantly present" and with regards to *Phascolosoma agassizi*, "undoubtedly many of these which were present in the area were overlooked".

In the Discussion, Provin notes the difficulty in attributing population changes with methodological differences when comparing his data to Hewatt's data. For example: "Though the data for this area [quadrat 35] shows a considerable difference between the two studies, I suspect that a considerable part of this difference is due to differences in methods rather than real differences in the populations existing on the area; specifically Hewatt probably removed the sand and algae from the rocks and examined them, while this was not done in the present study." Difficulties with species identification of similar of congeners were also discussed. In summary, "Little grounds were seen for trying to account for the observed population differences…more careful standardization of the procedure and a more stable taxonomy will be necessary for … determining actual population changes".

<u>Description of methods, 1993-1996 (Sagarin, Barry, Gilman, Baxter; SBGB)</u>

The following descriptions of the second resurvey of Hewatt's transect, initiated by R. Sagarin and S. Gilman under the guidance of C. Baxter, were paraphrased from a detailed manuscript by Sagarin, Barry, Gilman, and Baxter; hereafter referred to as SBGB (Sagarin et al., 1999). No personal interpretations were included unless explicitly noted.

Hewatt's transect was relocated with the aid of two of Hewatt's four original brass bolts that marked the transect. SBGB installed four 1.3cm diameter titanium plugs to mark the eastern edge of the transect. Sagarin and Gilman relocated and resurveyed 57 plots between spring 1993 and summer 1995. During summer 1996 SBGB resurveyed the first 19 plots surveyed in spring 1993; this included quadrats 27-38 and 62-68 (hereafter referred to as 'core' quadrats).

Counts were performed as nondestructively as possible, but some animals were removed temporarily to aid counting or identification. With few exceptions, all individuals within a plot were counted, including those on or under marine plants or other species. Several species were ignored deliberately. Species that could not be readily and nondestructively identified were not counted; only species that could be identified with the unaided eye were counted. Small animals that live abundantly in algal holdfasts (e.g., the bivalve *Lasea*, and gastropods *Barleeia*, *Caecum*, *Tricolia*) were ignored. SBGB note that "no such species were included in Hewatt's study, despite the presence of turf algae in Hewatt's photographs, and thus we assume that he also ignored them." Colonial organisms such as tunicates and sponges were not recorded, except those countable in discrete units (e.g., *Clavelina huntsmani*, *Polyclinum planum*). All species were counted directly, except for the barnacles *Balanus glandula* and *Chthamalus*; these were subsampled using 25 haphazard throws of a 2x2 and 3x3cm quadrat, respectively. Due to the high reported barnacle densities, SBGB note that Hewatt "very likely estimated barnacle abundance, although no such method is reported". Regional guides were used for species identification (Smith & Carlton, 1975; Morris, Abbott & Haderlie, 1980).

Description of methods, 1999 – 2014 (Sagarin)

During this period, R. Sagarin returned to Hopkins Marine Station to resurvey the core quadrats 27-38 and 62-68 during these years: 1999, 2002, 2005, 2009, 2014, 2015. No description of methods is available. We assume that the methods were similar to the methods described by SBGB for the years 1993-1996.

Description of methods, 2016-2019 (Watanabe, Micheli)

J. Watanabe and F. Micheli continued surveying the core quadrats on an annual basis after R. Sagarin's death in 2015. Surveys were conducted between May and July; the sampling effort often included students from an undergraduate spring course at HMS. R. Elahi began leading the survey effort in 2019, with transfer of knowledge from J. Watanabe and F. Micheli. In general, the sampling methods described by SBGB (Sagarin et al., 1999) were maintained.

Description of methods, 2020-2024 (Elahi)

R. Elahi and F. Micheli continued surveying the core quadrats, and started surveying quadrats 12, 16, 20, and 24 to address biodiversity change in the upper intertidal. In addition to the annual late spring sampling, a subset of quadrats was also sampled during winter months. The sampling methods remained the same, with a few exceptions. To count barnacles (*Balanus* and *Chthamalus*), one 10x10cm quadrat was placed haphazardly in each quarter of the quadrat (instead of 25 haphazard throws of 2x2 or 3x3cm quadrats). New species continued to be added to the database, and voucher specimens were collected starting in 2022. In addition, when species could not be identified to species (either in the field or lab), individuals were recorded to the lowest possible taxon. Amphipods and spirorbids were no longer counted in the quadrats. Regional guides were used for species identification (Morris, Abbott & Haderlie, 1980; Carlton, 2007). In general, 45 minutes to two hours were spent sampling each quadrat in the field. Mobile species that could be easily removed were brought into the lab for identification when necessary. Sand and shell hash were combed through to a depth of 5 to 10 cm (finger length). Living biogenic habitat (e.g., holdfasts, mussels) were not destructively sampled.

Description of methods, 2024 (Elahi)

R. Elahi surveyed the core quadrats and the extra quadrats (12, 16, 20, 24), with one modification. Hitchhikers on snails (e.g., *Crepidula*, *Lottia*) were no longer counted because this required a considerable amount of time in the field. In addition, as part of a collaboration testing eDNA methods and visual surveys (in collaboration with M. Shea), we sampled quadrats 21, 22, and 23.
