## Appendix 2 for "Passing the quadrat: inferring biodiversity change over time and across investigators"

### Appendix S2

#### Section S1. Supporting Tables

Table S1. Table of species names used in this paper, by Sagarin and coauthors (Barry et al. 1995, Sagarin et al. 1999), and by Hewatt (Hewatt 1937). The 'Analysis' column indicates the taxon used in this study for analyses of biodiversity change. NA indicates that the taxon was not recorded.

| Phylum | Taxon | Taxon (Sagarin) | Taxon (Hewatt) | Analysis |
|---|---|---|---|---|
| Annelida | Arabella iricolor | Arabella iricolor | NA | species |
| Annelida | Cirriformia spirabrancha | NA | NA | species |
| Annelida | Drilonereis spp | NA | NA | genus |
| Annelida | Errantia clade | NA | NA | subclass |
| Annelida | Glycera americana | Glycera americana | NA | species |
| Annelida | Halosydna brevisetosa | Halosydna insignis | Halosydna insignis | species |
| Annelida | Hyboscolex pacificus | NA | NA | species |
| Annelida | Leitoscoloplos pugettensis | NA | NA | species |
| Annelida | Lumbrineridae clade | NA | NA | family |
| Annelida | Lumbrineridae spA | NA | NA | family |
| Annelida | Lumbrineridae spB | NA | NA | family |
| Annelida | Lumbrineris spp | Lumbrinereis spp | Lumbrinereis sp | family |
| Annelida | Marphysa sanguinea | NA | NA | species |
| Annelida | Nereis spp | Nereis spp | Nereis sp | genus |
| Annelida | Nereis vexillosa | NA | NA | genus |
| Annelida | Oxydromus pugettensis | NA | NA | species |
| Annelida | Phascolosoma agassizii | Phascolosoma agassizii | Physcasoma agassizi | species |
| Annelida | Sedentaria clade | NA | NA | subclass |
| Annelida | Serpula columbiana | NA | Serpula columbiana | genus |
| Annelida | Serpula spp | NA | NA | genus |

(continued)

| Phylum | Taxon | Taxon (Sagarin) | Taxon (Hewatt) | Analysis |
|---|---|---|---|---|
| Annelida | Serpula vermicularis | NA | NA | genus |
| Annelida | Serpulidae clade | NA | NA | subclass |
| Arthropoda | Amphipoda clade | NA | NA | order |
| Arthropoda | Balanomorpha clade | NA | NA | order |
| Arthropoda | Balanus glandula | Balanus glandula | Balanus glandula | species |
| Arthropoda | Betaeus longidactylus | NA | NA | species |
| Arthropoda | Brachyura clade | NA | NA | class |
| Arthropoda | Cancer jordani | Cancer jordani | Cancer jordani | species |
| Arthropoda | Cancer productus | Cancer productus | Cancer productus | species |
| Arthropoda | Cancridae clade | NA | NA | family |
| Arthropoda | Chthamalus spp | Chthamalus spp | NA | genus |
| Arthropoda | Cirolana harfordi | Cirolana harfordi | Cirolana harfordi | species |
| Arthropoda | Cryptolithodes sitchensis | Cryptolithodes sitchensis | NA | species |
| Arthropoda | Hemigrapsus nudus | Hemigrapsus nudus | Hemigrapsus nudus | species |
| Arthropoda | Heptacarpus sitchensis | Heptacarpus pictus | Hippolyte californiensis | species |
| Arthropoda | Idotea spp | NA | NA | genus |
| Arthropoda | Idotea urotoma | Idotea urotoma | Idothea rectilinea | genus |
| Arthropoda | Isopoda clade | NA | NA | order |
| Arthropoda | Ligia occidentialis | Ligia occidentialis | Ligyda occidentalis | genus |
| Arthropoda | Lophopanopeus bellus | NA | NA | genus |
| Arthropoda | Lophopanopeus heathii | Lophopanopeus heathii | Lophopanopeus heathii | genus |
| Arthropoda | Lophopanopeus leucomanus | Lophopanopeus leucomanus | NA | genus |
| Arthropoda | Lophopanopeus spp | NA | NA | genus |
| Arthropoda | Loxorhynchus crispatus | Loxorhynchus crispatus | NA | species |

(continued)

| Phylum | Taxon | Taxon (Sagarin) | Taxon (Hewatt) | Analysis |
|---|---|---|---|---|
| Arthropoda | Megabalanus californicus | Megabalanus californicus | Balanus tintinnabulum californicus | species |
| Arthropoda | Pachycheles rudis | Pachycheles rudis | Pachycheles rudis | species |
| Arthropoda | Pachygrapsus crassipes | Pachygrapsus crassipes | Pachygrapsus crassipes | species |
| Arthropoda | Pagurus spp | Pagurus spp | NA | genus |
| Arthropoda | Paraxanthias taylori | Paraxanthias taylori | Xanthias taylori | species |
| Arthropoda | Petrolisthes cinctipes | Petrolisthes cinctipes | Petrolisthes cinctipes | genus |
| Arthropoda | Petrolisthes eriomerus | NA | NA | genus |
| Arthropoda | Petrolisthes spp | NA | NA | genus |
| Arthropoda | Pinnotheridae clade | NA | NA | family |
| Arthropoda | Pollicipes polymerus | Pollicipes polymerus | Mitella polymerus | species |
| Arthropoda | Pugettia foliata | Mimulus foliatus | Mimulus foliatus | species |
| Arthropoda | Pugettia producta | Pugettia producta | Pugettia productus | species |
| Arthropoda | Pugettia richii | Pugettia richii | Pugettia richii | species |
| Arthropoda | Pugettia spp | NA | NA | genus |
| Arthropoda | Pycnogonida clade | NA | NA | family |
| Arthropoda | Pycnogonum stearnsi | NA | NA | family |
| Arthropoda | Romaleon antennarium | Cancer antennarius | Cancer antennarius | species |
| Arthropoda | Spirontocaris picta | Spirontocaris picta | Spirontocaris picta | species |
| Arthropoda | Tetraclita rubescens | Tetraclita rubescens | Tetraclita squamosa rubescens | species |
| Bryozoa | Bryozoa clade | NA | NA | phylum |
| Bryozoa | Bugula neritina | NA | NA | species |
| Bryozoa | Cryptosula pallasiana | Cryptosula pallasiana | Hippodiplosia poliasiana | species |
| Bryozoa | Integripelta bilabiata | NA | NA | species |

(continued)

| Phylum | Taxon | Taxon (Sagarin) | Taxon (Hewatt) | Analysis |
|---|---|---|---|---|
| Bryozoa | Watersipora subtorquata | NA | NA | species |
| Chordata | Anoplarchus purpurescens | NA | NA | species |
| Chordata | Aplidium californicum | Aplidium californicum | Amaroucium californicum | genus |
| Chordata | Aplidium solidum | NA | NA | genus |
| Chordata | Aplidium spp | NA | NA | genus |
| Chordata | Ascidiacea clade | NA | NA | class |
| Chordata | Clavelina huntsmani | Clavelina huntsmani | Clavelina sp | species |
| Chordata | Distaplia occidentalis | NA | NA | species |
| Chordata | Eudistoma diaphanes | NA | NA | genus |
| Chordata | Eudistoma molle | Archidistoma molle | NA | genus |
| Chordata | Eudistoma psammion | NA | NA | genus |
| Chordata | Gibbonsia montereyensis | NA | NA | species |
| Chordata | Henricia leviuscula | Henricia leviuscula | NA | species |
| Chordata | Leptasterias spp | Leptasterias spp | Leptasterias aequalis | genus |
| Chordata | Oligocottus maculosus | Oligocottus maculosus | NA | species |
| Chordata | Perophora annectens | NA | NA | species |
| Chordata | Polyclinum planum | Polyclinum planum | Glossophorum planum | species |
| Chordata | Pycnoclavella stanleyi | NA | NA | species |
| Chordata | Rimicola eigenmanni | Rimicola eigenmanni | NA | species |
| Chordata | Synoicum spp | NA | NA | genus |
| Cnidaria | Abietinaria spp | NA | NA | genus |
| Cnidaria | Aglaophenia spp | NA | NA | genus |
| Cnidaria | Aglaophenia struthionides | NA | Aglaophenia struthionides | genus |
| Cnidaria | Anthopleura artemisia | NA | NA | genus |

(continued)

| Phylum | Taxon | Taxon (Sagarin) | Taxon (Hewatt) | Analysis |
|---|---|---|---|---|
| Cnidaria | Anthopleura elegantissima | Anthopleura elegantissima | Cribrina elegantissima | genus |
| Cnidaria | Anthopleura sola | Anthopleura sola | NA | genus |
| Cnidaria | Anthopleura xanthogrammica | Anthopleura xanthogrammica | Cribrina xanthogrammica | genus |
| Cnidaria | Balanophyllia elegans | Balanophyllia elegans | NA | species |
| Cnidaria | Corynactis californica | Corynactis californica | NA | species |
| Cnidaria | Epiactis prolifera | Epiactis prolifera | NA | species |
| Cnidaria | Eudendrium spp | NA | NA | genus |
| Cnidaria | Symplectoscyphus spp | NA | Sertularia pulchella | genus |
| Echinodermata | Amphiodia occidentalis | Amphiodia occidentalis | Amphiodia occidentalis | species |
| Echinodermata | Amphipholis pugetana | Amphipholis pugetana | Amphipholis pugetana | genus |
| Echinodermata | Amphipholis spp | NA | NA | genus |
| Echinodermata | Leptosynapta albicans | NA | NA | genus |
| Echinodermata | Leptosynapta inhaerens | Leptosynapta inhaerens | Leptosynapta inhaerens | genus |
| Echinodermata | Lissothuria nutriens | Lissothuria nutriens | Thyonepsolus nutriens | species |
| Echinodermata | Ophiactis simplex | Ophiactis simplex | NA | species |
| Echinodermata | Ophioderma panamensis | Ophioderma panamense | NA | species |
| Echinodermata | Ophiothrix spiculata | Ophiothrix spiculata | Ophiothrix spiculata | species |
| Echinodermata | Ophiuroidea clade | NA | NA | class |
| Echinodermata | Patiria miniata | Asterina miniata | Patiria miniata | species |
| Echinodermata | Pisaster ochraceus | Pisaster ochraceus | Pisaster ochraceus | species |
| Echinodermata | Strongylocentrotus purpuratus | Strongylocentrotus purpuratus | Strongylocentrotus purpuratus | species |
| Mollusca | Acanthinucella punctulata | Acanthina punctulata | Acanthina lapilloides | species |

(continued)

| Phylum | Taxon | Taxon (Sagarin) | Taxon (Hewatt) | Analysis |
|---|---|---|---|---|
| Mollusca | Acmaea mitra | Lottia mitra | Acmaea mitra | species |
| Mollusca | Alia carinata | Mitrella carinata | Columbella carinata | species |
| Mollusca | Amphissa columbiana | Amphissa columbiana | NA | genus |
| Mollusca | Amphissa versicolor | Amphissa versicolor | Amphissa versicolor | genus |
| Mollusca | Antisabia panamensis | Hipponix cranioides | Hipponix antiquatus | species |
| Mollusca | Aplysia californica | NA | NA | species |
| Mollusca | Atrimitra idae | NA | NA | species |
| Mollusca | Bivalvia clade | NA | NA | class |
| Mollusca | Californiconus californicus | NA | NA | species |
| Mollusca | Calliostoma annulatum | NA | NA | species |
| Mollusca | Calliostoma canaliculatum | Calliostoma canaliculatum | NA | genus |
| Mollusca | Calliostoma ligatum | Calliostoma ligatum | Calliostoma costatum | genus |
| Mollusca | Calliostoma spp | NA | NA | genus |
| Mollusca | Ceratostoma foliatum | Ceratostoma foliatum | NA | species |
| Mollusca | Chaetopleura gemma | Chaetopleura gemma | NA | species |
| Mollusca | Chama pellucida | Chama pellucida | Chama pellucida | species |
| Mollusca | Chlamys hastata | NA | NA | species |
| Mollusca | Coryphella spp | NA | NA | genus |
| Mollusca | Crassadoma gigantea | Hinnites giganteus | NA | species |
| Mollusca | Crepidula adunca | Crepidula adunca | Crepidula adunca | species |
| Mollusca | Cyanoplax hartwegii | Cyanoplax hartwegii | Lepidochitona hartwegii | species |
| Mollusca | Diaphoreolis lagunae | Cuthona lagunae | NA | species |
| Mollusca | Diaulula sandiegensis | NA | NA | species |
| Mollusca | Diodora aspera | NA | NA | species |
| Mollusca | Dirona spp | NA | NA | genus |

(continued)

| Phylum | Taxon | Taxon (Sagarin) | Taxon (Hewatt) | Analysis |
|---|---|---|---|---|
| Mollusca | Doridina clade | NA | NA | order |
| Mollusca | Doriopsilla albopunctata | Doriopsilla albopunctata | NA | species |
| Mollusca | Epitonium indianorum | NA | NA | genus |
| Mollusca | Epitonium tinctum | Epitonium tinctum | NA | genus |
| Mollusca | Eulithidium pulloides | NA | NA | genus |
| Mollusca | Fissurella volcano | Fissurella volcano | Fissurella volcano | species |
| Mollusca | Fissurellidea bimaculata | Megatebennus bimaculatus | NA | species |
| Mollusca | Flabellina trilineata | Coryphella trilineata | NA | species |
| Mollusca | Gari californica | NA | NA | species |
| Mollusca | Gastropoda clade | NA | NA | class |
| Mollusca | Geitodoris heathi | NA | Discodoris heathi | species |
| Mollusca | Gigahomalopoma luridum | Homalopoma luridum | Leptothyra carpenteri | species |
| Mollusca | Haliotis cracherodii | Haliotis cracherodii | NA | species |
| Mollusca | Hermissenda crassicornis | Hermissenda crassicornis | Hermissenda crassicornis | genus |
| Mollusca | Hermissenda opalescens | NA | NA | genus |
| Mollusca | Hesperaptyxis luteopictus | Fusinus luteopictus | NA | species |
| Mollusca | Hespererato vitellina | Erato vitellina | NA | species |
| Mollusca | Kelletia kelletii | NA | NA | species |
| Mollusca | Kellia laperousii | Kellia laperousii | Kellia laperousii | species |
| Mollusca | Lacuna marmorata | Lacuna marmorata | NA | genus |
| Mollusca | Lacuna porrecta | NA | NA | genus |
| Mollusca | Lacuna unifasciata | NA | NA | genus |
| Mollusca | Lepidozona cooperi | NA | NA | genus |

(continued)

| Phylum | Taxon | Taxon (Sagarin) | Taxon (Hewatt) | Analysis |
|---|---|---|---|---|
| Mollusca | Lepidozona mertensii | Lepidozona mertensii | NA | genus |
| Mollusca | Limacia cockerelli | Laila cockerelli | Laila cockerelli | species |
| Mollusca | Lirobittium spp | NA | NA | genus |
| Mollusca | Littorina keenae | Littorina keenae | Littorina planaxis | species |
| Mollusca | Littorina scutulata | Littorina scutulata | Littorina scutulata | species |
| Mollusca | Lottia alaska | NA | NA | genus |
| Mollusca | Lottia asmi | Lottia asmi | Acmaea asmi | species |
| Mollusca | Lottia digitalis | Lottia digitalis | Acmaea digitalis | species |
| Mollusca | Lottia fenestrata | NA | NA | genus |
| Mollusca | Lottia instabilis | NA | NA | genus |
| Mollusca | Lottia limatula | Lottia limatula | Acmaea limatula | species |
| Mollusca | Lottia ochracea | Lottia ochracea | NA | genus |
| Mollusca | Lottia paradigitalis | Lottia paradigitalis | NA | species |
| Mollusca | Lottia pelta | Lottia pelta | Acmaea cassis pelta hybrid | genus |
| Mollusca | Lottia persona | NA | NA | genus |
| Mollusca | Lottia scabra | Macclintockia scabra | Acmaea scabra | species |
| Mollusca | Lottia scutum | Tectura scutum | Acmaea patina | species |
| Mollusca | Lottia spp | NA | NA | genus |
| Mollusca | Margarites salmoneus | NA | NA | species |
| Mollusca | Mitrella tuberosa | NA | NA | species |
| Mollusca | Modiolus carpenteri | Modiolus carpenteri | NA | genus |
| Mollusca | Mopalia ciliata | NA | NA | genus |
| Mollusca | Mopalia lignosa | NA | NA | genus |
| Mollusca | Mopalia muscosa | Mopalia muscosa | Mopalia muscosa | genus |
| Mollusca | Mopalia spp | NA | NA | genus |
| Mollusca | Muricidae clade | NA | NA | family |
| Mollusca | Mytilus californianus | Mytilus californianus | Mytilus californianus | genus |
| Mollusca | Mytilus edulis | Mytilus edulis | NA | genus |

(continued)

| Phylum | Taxon | Taxon (Sagarin) | Taxon (Hewatt) | Analysis |
|---|---|---|---|---|
| Mollusca | Nassarius mendicus | Nassarius mendicus | NA | species |
| Mollusca | Neostylidium eschrichtii | Bittium eschrichtii | NA | species |
| Mollusca | Nucella emarginata | Nucella emarginata | Thais emarginata | genus |
| Mollusca | Nucella lamellosa | NA | NA | genus |
| Mollusca | Nucella spp | NA | NA | genus |
| Mollusca | Nudibranchia clade | NA | NA | order |
| Mollusca | Nuttallina californica | Nuttallina californica | Nuttallina californica | species |
| Mollusca | Odetta fetella | NA | NA | species |
| Mollusca | Okenia rosacea | Hopkinsia rosacea | Hopkinsia rosacea | species |
| Mollusca | Onchidella carpenteri | Onchidella borealis | NA | species |
| Mollusca | Paciocinebrina atropurpurea | NA | NA | genus |
| Mollusca | Paciocinebrina circumtexta | Ocenebra circumtexta | NA | species |
| Mollusca | Paciocinebrina interfossa | NA | Tritonalia interfossa | genus |
| Mollusca | Paciocinebrina lurida | Ocenebra lurida | Tritonalia lurida | genus |
| Mollusca | Paciocinebrina spp | NA | NA | genus |
| Mollusca | Paciocinebrina subangulata | NA | NA | genus |
| Mollusca | Petaloconchus montereyensis | NA | NA | species |
| Mollusca | Polyplacophora clade | NA | NA | class |
| Mollusca | Pseudochama exogyra | NA | NA | species |
| Mollusca | Pseudomelatoma torosa | Pseudomelatoma torosa | NA | species |
| Mollusca | Pseudopusula californiana | NA | NA | species |

(continued)

| Phylum | Taxon | Taxon (Sagarin) | Taxon (Hewatt) | Analysis |
|---|---|---|---|---|
| Mollusca | Rostanga pulchra | Rostanga pulchra | Rostanga pulchra | species |
| Mollusca | Tectura paleacea | Tectura paleacea | NA | species |
| Mollusca | Tegula brunnea | Tegula brunnea | Tegula brunnea | species |
| Mollusca | Tegula funebralis | Tegula funebralis | Tegula funebralis | species |
| Mollusca | Tegula montereyi | Tegula montereyi | NA | species |
| Mollusca | Tegula pulligo | Tegula pulligo | NA | species |
| Mollusca | Thylacodes squamigerus | Serpulorbis squamigerus | NA | species |
| Mollusca | Tonicella lineata | Tonicella lineata | Lepidochitona lineata | species |
| Mollusca | Triopha catalinae | NA | Triopha carpenteri | species |
| Mollusca | Urosalpinx cinerea | NA | NA | species |
| Nemertea | Emplectonema gracile | Emplectonema gracile | NA | species |
| Nemertea | Nemertea clade | NA | NA | phylum |
| Nemertea | Paranemertes peregrina | Paranemertes peregrina | Paranemertes peregrina | species |
| Platyhelminthes | Notocomplana acticola | NA | NA | genus |
| Platyhelminthes | Notocomplana spp | NA | NA | genus |
| Platyhelminthes | Platyhelminthes clade | NA | NA | phylum |
| Platyhelminthes | Pseudoalloioplana californica | Alloioplana californica | Planocera californica | species |
| Porifera | Antho karykina | Plocamia karykina | Plocamia karykinos | species |
| Porifera | Clathria pennata | NA | NA | species |
| Porifera | Haliclona spp | NA | NA | genus |
| Porifera | Porifera clade | NA | NA | phylum |
| Sipuncula | Sipuncula clade | NA | NA | order |

Table S2. Complete list of taxa observed (TRUE) or not observed (FALSE) in each investigator era.

| Taxon | 1931-1933 | 1949 | 1993,1996 | 1999-2015 | 2016-2019 | 2020-2023 |
|---|---|---|---|---|---|---|
| Abietinaria spp | FALSE | TRUE | FALSE | FALSE | TRUE | TRUE |
| Acanthinucella punctulata | TRUE | TRUE | TRUE | TRUE | TRUE | TRUE |
| Acmaea mitra | TRUE | FALSE | TRUE | TRUE | TRUE | TRUE |
| Aglaophenia spp | FALSE | FALSE | FALSE | FALSE | TRUE | TRUE |
| Aglaophenia struthionides | FALSE | TRUE | FALSE | FALSE | FALSE | FALSE |
| Alia carinata | TRUE | FALSE | TRUE | TRUE | TRUE | TRUE |
| Amphiodia occidentalis | FALSE | FALSE | FALSE | FALSE | FALSE | TRUE |
| Amphipholis pugetana | TRUE | FALSE | FALSE | FALSE | FALSE | FALSE |
| Amphipholis spp | FALSE | FALSE | FALSE | FALSE | TRUE | FALSE |
| Amphissa columbiana | FALSE | FALSE | TRUE | FALSE | FALSE | FALSE |
| Amphissa versicolor | TRUE | TRUE | TRUE | TRUE | TRUE | TRUE |
| Anoplarchus purpurescens | FALSE | FALSE | FALSE | FALSE | FALSE | TRUE |
| Antho karykina | TRUE | FALSE | FALSE | FALSE | FALSE | FALSE |
| Anthopleura artemisia | FALSE | FALSE | TRUE | FALSE | TRUE | FALSE |
| Anthopleura elegantissima | TRUE | TRUE | FALSE | FALSE | TRUE | TRUE |
| Anthopleura sola | FALSE | FALSE | TRUE | TRUE | TRUE | TRUE |
| Anthopleura xanthogrammica | TRUE | TRUE | TRUE | TRUE | FALSE | TRUE |
| Antisabia panamensis | FALSE | FALSE | FALSE | TRUE | FALSE | FALSE |
| Aplidium californicum | TRUE | FALSE | TRUE | FALSE | TRUE | TRUE |
| Aplidium solidum | FALSE | FALSE | FALSE | FALSE | FALSE | TRUE |
| Aplysia californica | FALSE | FALSE | FALSE | FALSE | FALSE | TRUE |
| Arabella iricolor | FALSE | FALSE | TRUE | FALSE | FALSE | FALSE |
| Atrimitra idae | FALSE | FALSE | FALSE | FALSE | FALSE | TRUE |
| Balanophyllia elegans | FALSE | FALSE | FALSE | FALSE | FALSE | FALSE |
| Balanus glandula | TRUE | TRUE | TRUE | TRUE | TRUE | TRUE |
| Betaeus longidactylus | FALSE | TRUE | FALSE | FALSE | FALSE | FALSE |
| Bugula neritina | TRUE | TRUE | FALSE | FALSE | TRUE | TRUE |

(continued)

| Taxon | 1931-1933 | 1949 | 1993,1996 | 1999-2015 | 2016-2019 | 2020-2023 |
|---|---|---|---|---|---|---|
| Californiconus californicus | FALSE | FALSE | FALSE | FALSE | TRUE | TRUE |
| Calliostoma annulatum | FALSE | FALSE | TRUE | TRUE | FALSE | TRUE |
| Calliostoma canaliculatum | FALSE | FALSE | TRUE | TRUE | TRUE | TRUE |
| Calliostoma ligatum | TRUE | FALSE | TRUE | TRUE | TRUE | TRUE |
| Cancer jordani | FALSE | FALSE | FALSE | FALSE | FALSE | TRUE |
| Cancer productus | TRUE | TRUE | FALSE | FALSE | TRUE | FALSE |
| Ceratostoma foliatum | TRUE | TRUE | TRUE | TRUE | TRUE | TRUE |
| Chaetopleura gemma | TRUE | TRUE | TRUE | FALSE | FALSE | FALSE |
| Chama pellucida | TRUE | FALSE | FALSE | FALSE | FALSE | FALSE |
| Chlamys hastata | FALSE | FALSE | FALSE | FALSE | FALSE | TRUE |
| Chthamalus spp | FALSE | FALSE | TRUE | TRUE | TRUE | TRUE |
| Cirolana harfordi | TRUE | TRUE | FALSE | FALSE | TRUE | TRUE |
| Cirriformia spirabrancha | FALSE | FALSE | FALSE | FALSE | FALSE | TRUE |
| Clathria pennata | TRUE | TRUE | FALSE | FALSE | FALSE | FALSE |
| Clavelina huntsmani | TRUE | FALSE | TRUE | TRUE | TRUE | TRUE |
| Corynactis californica | FALSE | FALSE | TRUE | TRUE | FALSE | FALSE |
| Coryphella spp | FALSE | FALSE | FALSE | FALSE | TRUE | FALSE |
| Crassadoma gigantea | TRUE | FALSE | FALSE | FALSE | TRUE | FALSE |
| Crepidula adunca | TRUE | FALSE | TRUE | TRUE | TRUE | TRUE |
| Cryptolithodes sitchensis | FALSE | FALSE | FALSE | FALSE | TRUE | FALSE |
| Cryptosula pallasiana | TRUE | FALSE | FALSE | FALSE | FALSE | FALSE |
| Cyanoplax hartwegii | TRUE | FALSE | TRUE | TRUE | TRUE | TRUE |
| Diaphoreolis lagunae | FALSE | FALSE | TRUE | FALSE | FALSE | FALSE |
| Diaulula sandiegensis | TRUE | FALSE | FALSE | FALSE | FALSE | TRUE |
| Diodora aspera | TRUE | FALSE | FALSE | TRUE | FALSE | FALSE |
| Dirona spp | TRUE | FALSE | FALSE | FALSE | TRUE | FALSE |
| Distaplia occidentalis | FALSE | FALSE | FALSE | FALSE | TRUE | FALSE |
| Doriopsilla albopunctata | TRUE | FALSE | FALSE | FALSE | TRUE | FALSE |

(continued)

| Taxon | 1931-1933 | 1949 | 1993,1996 | 1999-2015 | 2016-2019 | 2020-2023 |
|---|---|---|---|---|---|---|
| Drilonereis spp | FALSE | TRUE | FALSE | FALSE | FALSE | TRUE |
| Emplectonema gracile | FALSE | FALSE | TRUE | FALSE | FALSE | FALSE |
| Epiactis prolifera | TRUE | FALSE | TRUE | TRUE | TRUE | FALSE |
| Epitonium indianorum | FALSE | FALSE | FALSE | FALSE | TRUE | FALSE |
| Epitonium tinctum | FALSE | FALSE | TRUE | FALSE | TRUE | TRUE |
| Eudendrium spp | FALSE | FALSE | FALSE | FALSE | TRUE | FALSE |
| Eudistoma diaphanes | FALSE | FALSE | FALSE | FALSE | FALSE | TRUE |
| Eudistoma molle | FALSE | FALSE | TRUE | FALSE | TRUE | TRUE |
| Eudistoma psammion | FALSE | FALSE | FALSE | FALSE | TRUE | TRUE |
| Eulithidium pulloides | FALSE | FALSE | FALSE | FALSE | TRUE | FALSE |
| Fissurella volcano | TRUE | FALSE | TRUE | TRUE | TRUE | TRUE |
| Fissurellidea bimaculata | TRUE | FALSE | FALSE | TRUE | FALSE | TRUE |
| Flabellina trilineata | FALSE | FALSE | TRUE | FALSE | FALSE | FALSE |
| Gari californica | FALSE | FALSE | FALSE | FALSE | FALSE | TRUE |
| Geitodoris heathi | TRUE | FALSE | FALSE | FALSE | TRUE | FALSE |
| Gibbonsia montereyensis | FALSE | FALSE | FALSE | FALSE | FALSE | TRUE |
| Gigahomalopoma luridum | TRUE | FALSE | TRUE | TRUE | TRUE | TRUE |
| Glycera americana | FALSE | TRUE | TRUE | FALSE | FALSE | TRUE |
| Haliclona spp | FALSE | FALSE | FALSE | FALSE | FALSE | TRUE |
| Haliotis cracherodii | TRUE | FALSE | TRUE | FALSE | FALSE | TRUE |
| Halosydna brevisetosa | TRUE | FALSE | FALSE | FALSE | TRUE | TRUE |
| Hemigrapsus nudus | TRUE | FALSE | FALSE | FALSE | TRUE | TRUE |
| Henricia leviuscula | TRUE | TRUE | TRUE | FALSE | TRUE | TRUE |
| Heptacarpus sitchensis | TRUE | FALSE | FALSE | FALSE | TRUE | FALSE |
| Hermissenda crassicornis | FALSE | FALSE | TRUE | TRUE | TRUE | TRUE |
| Hermissenda opalescens | FALSE | FALSE | FALSE | FALSE | TRUE | TRUE |
| Hesperaptyxis luteopictus | FALSE | FALSE | TRUE | TRUE | TRUE | FALSE |
| Hespererato vitellina | FALSE | FALSE | TRUE | TRUE | TRUE | FALSE |

(continued)

| Taxon | 1931-1933 | 1949 | 1993,1996 | 1999-2015 | 2016-2019 | 2020-2023 |
|---|---|---|---|---|---|---|
| Hyboscolex pacificus | FALSE | TRUE | FALSE | FALSE | FALSE | TRUE |
| Idotea spp | FALSE | FALSE | FALSE | FALSE | FALSE | TRUE |
| Idotea urotoma | FALSE | TRUE | FALSE | FALSE | TRUE | FALSE |
| Integripelta bilabiata | FALSE | TRUE | FALSE | FALSE | TRUE | TRUE |
| Kelletia kelletii | FALSE | FALSE | FALSE | FALSE | FALSE | TRUE |
| Kellia laperousii | TRUE | FALSE | FALSE | FALSE | FALSE | FALSE |
| Lacuna marmorata | FALSE | FALSE | TRUE | TRUE | FALSE | TRUE |
| Lacuna porrecta | FALSE | FALSE | FALSE | FALSE | TRUE | FALSE |
| Lacuna unifasciata | FALSE | FALSE | FALSE | FALSE | FALSE | TRUE |
| Leitoscoloplos pugettensis | FALSE | FALSE | FALSE | FALSE | FALSE | TRUE |
| Lepidozona cooperi | FALSE | FALSE | FALSE | FALSE | TRUE | FALSE |
| Lepidozona mertensii | TRUE | TRUE | TRUE | FALSE | FALSE | FALSE |
| Leptasterias spp | TRUE | TRUE | TRUE | TRUE | TRUE | TRUE |
| Leptosynapta albicans | FALSE | FALSE | FALSE | FALSE | FALSE | TRUE |
| Leptosynapta inhaerens | TRUE | FALSE | FALSE | FALSE | FALSE | FALSE |
| Ligia occidentialis | TRUE | TRUE | FALSE | FALSE | FALSE | FALSE |
| Limacia cockerelli | FALSE | FALSE | FALSE | FALSE | TRUE | FALSE |
| Lirobittium spp | TRUE | TRUE | FALSE | FALSE | TRUE | FALSE |
| Lissothuria nutriens | FALSE | FALSE | FALSE | FALSE | FALSE | TRUE |
| Littorina keenae | TRUE | TRUE | TRUE | FALSE | FALSE | TRUE |
| Littorina scutulata | TRUE | TRUE | TRUE | TRUE | TRUE | TRUE |
| Lophopanopeus bellus | FALSE | FALSE | FALSE | FALSE | FALSE | TRUE |
| Lophopanopeus heathii | FALSE | TRUE | FALSE | FALSE | FALSE | FALSE |
| Lophopanopeus leucomanus | FALSE | FALSE | TRUE | FALSE | FALSE | FALSE |
| Lottia alaska | FALSE | FALSE | FALSE | FALSE | FALSE | TRUE |
| Lottia asmi | TRUE | TRUE | TRUE | TRUE | TRUE | TRUE |
| Lottia digitalis | TRUE | TRUE | TRUE | TRUE | TRUE | TRUE |
| Lottia fenestrata | FALSE | FALSE | FALSE | FALSE | FALSE | TRUE |

(continued)

| Taxon | 1931-1933 | 1949 | 1993,1996 | 1999-2015 | 2016-2019 | 2020-2023 |
|---|---|---|---|---|---|---|
| Lottia instabilis | FALSE | FALSE | FALSE | FALSE | TRUE | FALSE |
| Lottia limatula | TRUE | TRUE | TRUE | TRUE | TRUE | TRUE |
| Lottia ochracea | FALSE | FALSE | FALSE | FALSE | TRUE | FALSE |
| Lottia paradigitalis | FALSE | FALSE | TRUE | TRUE | TRUE | TRUE |
| Lottia pelta | TRUE | TRUE | TRUE | TRUE | TRUE | TRUE |
| Lottia persona | FALSE | FALSE | FALSE | FALSE | FALSE | TRUE |
| Lottia scabra | TRUE | TRUE | TRUE | TRUE | TRUE | TRUE |
| Lottia scutum | TRUE | TRUE | TRUE | TRUE | TRUE | TRUE |
| Loxorhynchus crispatus | FALSE | FALSE | FALSE | FALSE | TRUE | FALSE |
| Lumbrineridae spA | FALSE | FALSE | FALSE | FALSE | FALSE | TRUE |
| Lumbrineridae spB | FALSE | FALSE | FALSE | FALSE | FALSE | TRUE |
| Lumbrineris spp | TRUE | FALSE | FALSE | FALSE | FALSE | FALSE |
| Margarites salmoneus | FALSE | FALSE | FALSE | FALSE | FALSE | TRUE |
| Marphysa sanguinea | FALSE | FALSE | FALSE | FALSE | FALSE | TRUE |
| Megabalanus californicus | TRUE | FALSE | FALSE | FALSE | FALSE | FALSE |
| Mitrella tuberosa | FALSE | FALSE | FALSE | FALSE | TRUE | FALSE |
| Modiolus carpenteri | FALSE | FALSE | FALSE | FALSE | FALSE | TRUE |
| Mopalia ciliata | FALSE | FALSE | FALSE | FALSE | FALSE | TRUE |
| Mopalia lignosa | FALSE | FALSE | TRUE | FALSE | TRUE | TRUE |
| Mopalia muscosa | TRUE | FALSE | TRUE | TRUE | TRUE | TRUE |
| Mytilus californianus | TRUE | FALSE | TRUE | TRUE | TRUE | TRUE |
| Mytilus edulis | FALSE | FALSE | FALSE | FALSE | FALSE | TRUE |
| Nassarius mendicus | FALSE | FALSE | TRUE | FALSE | FALSE | FALSE |
| Neostylidium eschrichtii | TRUE | TRUE | TRUE | TRUE | TRUE | TRUE |
| Nereis spp | TRUE | FALSE | FALSE | FALSE | FALSE | TRUE |
| Notocomplana acticola | FALSE | TRUE | FALSE | FALSE | FALSE | TRUE |
| Nucella emarginata | FALSE | FALSE | FALSE | FALSE | TRUE | FALSE |
| Nucella lamellosa | FALSE | FALSE | FALSE | FALSE | TRUE | FALSE |
| Nuttallina californica | FALSE | FALSE | TRUE | TRUE | TRUE | TRUE |

(continued)

| Taxon | 1931-1933 | 1949 | 1993,1996 | 1999-2015 | 2016-2019 | 2020-2023 |
|---|---|---|---|---|---|---|
| Odetta fetella | FALSE | FALSE | FALSE | FALSE | FALSE | TRUE |
| Okenia rosacea | TRUE | FALSE | FALSE | TRUE | FALSE | FALSE |
| Oligocottus maculosus | TRUE | TRUE | TRUE | FALSE | FALSE | FALSE |
| Onchidella carpenteri | FALSE | FALSE | TRUE | FALSE | FALSE | FALSE |
| Ophiactis simplex | FALSE | FALSE | TRUE | FALSE | FALSE | FALSE |
| Ophioderma panamensis | FALSE | FALSE | TRUE | FALSE | FALSE | FALSE |
| Ophiothrix spiculata | FALSE | FALSE | TRUE | FALSE | TRUE | TRUE |
| Oxydromus pugettensis | FALSE | FALSE | FALSE | FALSE | FALSE | TRUE |
| Pachycheles rudis | TRUE | FALSE | FALSE | FALSE | FALSE | TRUE |
| Pachygrapsus crassipes | TRUE | TRUE | TRUE | TRUE | TRUE | TRUE |
| Paciocinebrina atropurpurea | FALSE | FALSE | FALSE | FALSE | TRUE | TRUE |
| Paciocinebrina circumtexta | FALSE | FALSE | TRUE | TRUE | TRUE | TRUE |
| Paciocinebrina interfossa | TRUE | FALSE | FALSE | FALSE | FALSE | TRUE |
| Paciocinebrina lurida | TRUE | FALSE | FALSE | FALSE | TRUE | TRUE |
| Paciocinebrina subangulata | FALSE | FALSE | FALSE | FALSE | TRUE | TRUE |
| Pagurus spp | TRUE | TRUE | TRUE | TRUE | TRUE | TRUE |
| Paranemertes peregrina | TRUE | TRUE | TRUE | FALSE | FALSE | TRUE |
| Paraxanthias taylori | FALSE | FALSE | TRUE | FALSE | FALSE | FALSE |
| Patiria miniata | FALSE | TRUE | TRUE | TRUE | TRUE | TRUE |
| Perophora annectens | FALSE | FALSE | FALSE | FALSE | FALSE | TRUE |
| Petaloconchus montereyensis | FALSE | FALSE | FALSE | TRUE | FALSE | FALSE |
| Petrolisthes cinctipes | TRUE | FALSE | FALSE | TRUE | TRUE | TRUE |
| Petrolisthes eriomerus | FALSE | FALSE | FALSE | FALSE | FALSE | TRUE |
| Phascolosoma agassizii | TRUE | FALSE | FALSE | FALSE | FALSE | TRUE |
| Pisaster ochraceus | TRUE | FALSE | TRUE | TRUE | TRUE | TRUE |
| Pollicipes polymerus | FALSE | FALSE | FALSE | FALSE | FALSE | TRUE |
| Polyclinum planum | TRUE | FALSE | TRUE | TRUE | TRUE | TRUE |
| Pseudoalloioplana californica | TRUE | TRUE | FALSE | FALSE | FALSE | FALSE |

(continued)

| Taxon | 1931-1933 | 1949 | 1993,1996 | 1999-2015 | 2016-2019 | 2020-2023 |
|---|---|---|---|---|---|---|
| Pseudochama exogyra | FALSE | FALSE | FALSE | FALSE | FALSE | TRUE |
| Pseudomelatoma torosa | TRUE | FALSE | TRUE | TRUE | TRUE | TRUE |
| Pseudopusula californiana | FALSE | FALSE | FALSE | FALSE | FALSE | TRUE |
| Pugettia foliata | FALSE | FALSE | FALSE | FALSE | FALSE | TRUE |
| Pugettia producta | TRUE | TRUE | TRUE | TRUE | TRUE | TRUE |
| Pugettia richii | FALSE | FALSE | TRUE | TRUE | TRUE | TRUE |
| Pycnoclavella stanleyi | FALSE | FALSE | FALSE | FALSE | TRUE | FALSE |
| Pycnogonum stearnsi | TRUE | FALSE | FALSE | FALSE | FALSE | TRUE |
| Rimicola eigenmanni | FALSE | FALSE | FALSE | FALSE | FALSE | TRUE |
| Romaleon antennarium | TRUE | TRUE | TRUE | FALSE | TRUE | TRUE |
| Rostanga pulchra | TRUE | FALSE | TRUE | TRUE | FALSE | TRUE |
| Serpula columbiana | TRUE | TRUE | FALSE | FALSE | TRUE | FALSE |
| Serpula vermicularis | FALSE | FALSE | TRUE | TRUE | FALSE | FALSE |
| Spirontocaris picta | FALSE | TRUE | FALSE | FALSE | FALSE | FALSE |
| Strongylocentrotus purpuratus | TRUE | TRUE | TRUE | TRUE | TRUE | TRUE |
| Symplectoscyphus spp | TRUE | TRUE | FALSE | FALSE | TRUE | FALSE |
| Synoicum spp | FALSE | FALSE | FALSE | FALSE | TRUE | TRUE |
| Tectura paleacea | FALSE | FALSE | TRUE | TRUE | TRUE | TRUE |
| Tegula brunnea | TRUE | FALSE | TRUE | TRUE | TRUE | TRUE |
| Tegula funebralis | TRUE | TRUE | TRUE | TRUE | TRUE | TRUE |
| Tegula montereyi | FALSE | FALSE | TRUE | FALSE | FALSE | TRUE |
| Tegula pulligo | FALSE | FALSE | TRUE | FALSE | TRUE | TRUE |
| Tetraclita rubescens | FALSE | FALSE | TRUE | TRUE | TRUE | TRUE |
| Thylacodes squamigerus | FALSE | FALSE | TRUE | TRUE | TRUE | TRUE |
| Tonicella lineata | TRUE | FALSE | TRUE | TRUE | TRUE | TRUE |
| Triopha catalinae | TRUE | TRUE | FALSE | FALSE | TRUE | FALSE |
| Urosalpinx cinerea | FALSE | FALSE | FALSE | FALSE | FALSE | TRUE |
| Watersipora subtorquata | FALSE | FALSE | FALSE | FALSE | FALSE | TRUE |

Table S3. List of processing steps and associated observations (i.e. total abundance) to prepare the quantitative dataset for analysis.

| Processing step | Observations | Proportion |
|---|---|---|
| All quantitative observations | 365914 | 1.00 |
| Removed amphipods and Spirorbis | 364895 | 1.00 |
| Removed limpets on Tegula | 362559 | 0.99 |
| Removed Pagurus | 347700 | 0.95 |
| Removed taxa above genus | 347543 | 0.95 |

Table S4. Total numbers of unique species, genera, clades, and taxa in the quantitative dataset, before and after data processing for analysis of temporal trends in biodiversity. Data processing included lumping species to genera and removing higher level taxa; see Methods for details.

| Quantitative data | Species | Genera | Clades | Taxa (total) |
|---|---|---|---|---|
| Pre-processed | 180 | 28 | 24 | 232 |
| Post-processed | 119 | 38 | 0 | 157 |

Table S5. Sampling effort for 23 unique quadrats on the permanent intertidal transect (1931-2023) initiated by Willis Hewatt, summarized by six distinct eras led by different primary investigators. 'Total quadrats' denotes the total number of quadrats that were sampled during each era (i.e., the unique quadrats may have been sampled more than once).

| Era | Sampling events | Investigator(s) | Unique quadrats | Total quadrats | Unique taxa | Total count | Total count (no balanoid barnacles) |
|---|---|---|---|---|---|---|---|
| 1931-1933 | 1 | Hewatt | 23 | 23 | 46 | 23263 | 9126 |
| 1949 | 1 | Provin | 3 | 3 | 27 | 18448 | 3148 |
| 1993,1996 | 2 | Sagarin, Barry, Gilman, Baxter | 19 | 38 | 74 | 30430 | 13365 |
| 1999-2015 | 7 | Sagarin | 19 | 133 | 55 | 49969 | 30412 |
| 2016-2019 | 4 | Micheli, Watanabe | 19 | 76 | 88 | 32496 | 32197 |
| 2020-2023 | 4 | Elahi | 23 | 118 | 105 | 155206 | 71121 |

Table S6. Spearman rank statistics for temporal trends (1931-2023) in diversity metrics.

| Metric | Scale | Rho | S statistic | P |
|---|---|---|---|---|
| Richness | quadrat | 0.310 | 668.845 | 0.211 |
| Hill-Shannon | quadrat | -0.692 | 1640.000 | 0.002 |
| Hill-Simpson | quadrat | -0.825 | 1768.000 | 0.000 |
| Richness | site | 0.247 | 730.000 | 0.322 |
| Hill-Shannon | site | -0.496 | 1450.000 | 0.038 |
| Hill-Simpson | site | -0.422 | 1378.000 | 0.082 |

Table S7. Average (SD) density (no. $m^{-2}$) for each genus by era. Standard deviation is not available for 1931-1933 because each quadrat was sampled only once. These 36 genera represent 99% of the individuals counted across all eras in the quantitative dataset.

| Phylum | Genus | 1931-1933 | 1993,1996 | 1999-2015 | 2016-2019 | 2020-2023 |
|---|---|---|---|---|---|---|
| Annelida | Halosydna | 4.0 | 0 (0) | 0 (0) | 0 (0) | 0.1 (0.2) |
| Annelida | Phascolosoma | 3.7 | 0 (0) | 0 (0) | 0 (0) | 0.1 (0.2) |
| Arthropoda | Balanus | 0.0 | 121.3 (167.1) | 0.3 (0.5) | 0.9 (0.7) | 0 (0) |
| Arthropoda | Chthamalus | 0.0 | 254.1 (63.8) | 122.6 (189.3) | 2.3 (3) | 129.8 (250.8) |
| Arthropoda | Cirolana | 0.0 | 0 (0) | 0 (0) | 0.1 (0.1) | 2 (3.8) |
| Arthropoda | Pachycheles | 8.5 | 0 (0) | 0 (0) | 0 (0) | 0.3 (0.5) |
| Arthropoda | Pachygrapsus | 2.1 | 0.9 (0.2) | 0.3 (0.3) | 0.6 (0.4) | 0.9 (0.1) |
| Arthropoda | Petrolisthes | 3.7 | 0 (0) | 0 (0.1) | 0.1 (0.1) | 0.8 (0.6) |
| Arthropoda | Pugettia | 1.8 | 0.7 (0.1) | 0.3 (0.1) | 0.2 (0.1) | 0.3 (0.2) |
| Arthropoda | Tetraclita | 0.0 | 22.3 (23.4) | 6.6 (10.2) | 8.4 (15) | 0 (0.1) |
| Chordata | Aplidium | 0.0 | 0 (0) | 0 (0) | 1 (0.7) | 0.9 (0.8) |
| Chordata | Clavelina | 17.3 | 12.4 (4.8) | 3.9 (4.1) | 16.6 (9.8) | 31.8 (50.6) |
| Chordata | Eudistoma | 0.0 | 0.2 (0.2) | 0 (0) | 0.1 (0.2) | 0.7 (0.9) |
| Chordata | Leptasterias | 1.9 | 0.8 (0.3) | 0.1 (0.1) | 0.1 (0) | 0.2 (0.1) |
| Cnidaria | Anthopleura | 1.5 | 5 (0.3) | 4.9 (0.7) | 5 (0.2) | 4.1 (1.1) |
| Cnidaria | Corynactis | 0.0 | 5.9 (0.1) | 4.6 (4.2) | 0 (0) | 0 (0) |
| Echinodermata | Amphipholis | 4.1 | 0 (0) | 0 (0) | 0 (0) | 0 (0) |
| Echinodermata | Strongylocentrotus | 11.3 | 1.3 (0.5) | 0.6 (0.7) | 0.9 (0.9) | 7.1 (5.6) |
| Mollusca | Acanthinucella | 0.0 | 0.5 (0.5) | 0.2 (0.1) | 0.2 (0.2) | 0.8 (0.5) |
| Mollusca | Acmaea | 0.6 | 0.1 (0.1) | 0.1 (0.1) | 0.2 (0.1) | 0.2 (0.4) |
| Mollusca | Alia | 122.4 | 5 (1.6) | 0.8 (0.5) | 0.6 (0.8) | 0.9 (1.1) |
| Mollusca | Amphissa | 9.1 | 2.2 (1.5) | 0.2 (0.1) | 0.6 (0.5) | 3 (3.5) |
| Mollusca | Calliostoma | 0.5 | 1.5 (0.9) | 1 (0.6) | 0.9 (0.4) | 0.4 (0.3) |
| Mollusca | Crepidula | 11.3 | 19.5 (5.4) | 19.6 (6.9) | 22.1 (10.5) | 20.6 (6.5) |
| Mollusca | Fissurella | 0.7 | 1.7 (0.5) | 0.4 (0.2) | 0.5 (0.3) | 0.4 (0.4) |
| Mollusca | Gigahomalopoma | 0.0 | 3.1 (4.2) | 0.1 (0.2) | 0.1 (0.2) | 0.2 (0.3) |

(continued)

| Phylum | Genus | 1931-1933 | 1993,1996 | 1999-2015 | 2016-2019 | 2020-2023 |
|---|---|---|---|---|---|---|
| Mollusca | Lacuna | 0.0 | 3.5 (1.6) | 1.5 (1.4) | 0 (0) | 0 (0) |
| Mollusca | Littorina | 0.0 | 0.4 (0.4) | 0.1 (0.2) | 0.1 (0.1) | 0.4 (0.2) |
| Mollusca | Lottia | 39.8 | 4.2 (2.1) | 4.5 (6.1) | 7.3 (6.3) | 3.8 (1.7) |
| Mollusca | Mopalia | 0.6 | 0.4 (0.1) | 0 (0) | 0.1 (0.1) | 0.5 (0.4) |
| Mollusca | Mytilus | 9.3 | 0.3 (0.2) | 0.2 (0.1) | 3.2 (2.7) | 87.2 (88.6) |
| Mollusca | Neostylidium | 0.0 | 4.4 (0.4) | 0.1 (0.1) | 0 (0) | 0.1 (0.1) |
| Mollusca | Nuttallina | 0.0 | 0.1 (0.2) | 0.2 (0.2) | 0.1 (0.2) | 0.1 (0.1) |
| Mollusca | Paciocinebrina | 0.2 | 1 (0.2) | 0.5 (0.3) | 0.8 (0.3) | 2.4 (0.8) |
| Mollusca | Tegula | 47.3 | 181.6 (13.4) | 132 (42.3) | 280.8 (30.2) | 294.7 (29.5) |
| Mollusca | Thylacodes | 0.0 | 10.9 (5) | 7.5 (5.1) | 1.6 (1) | 1.2 (0.9) |

#### Section S2. Supporting Figures

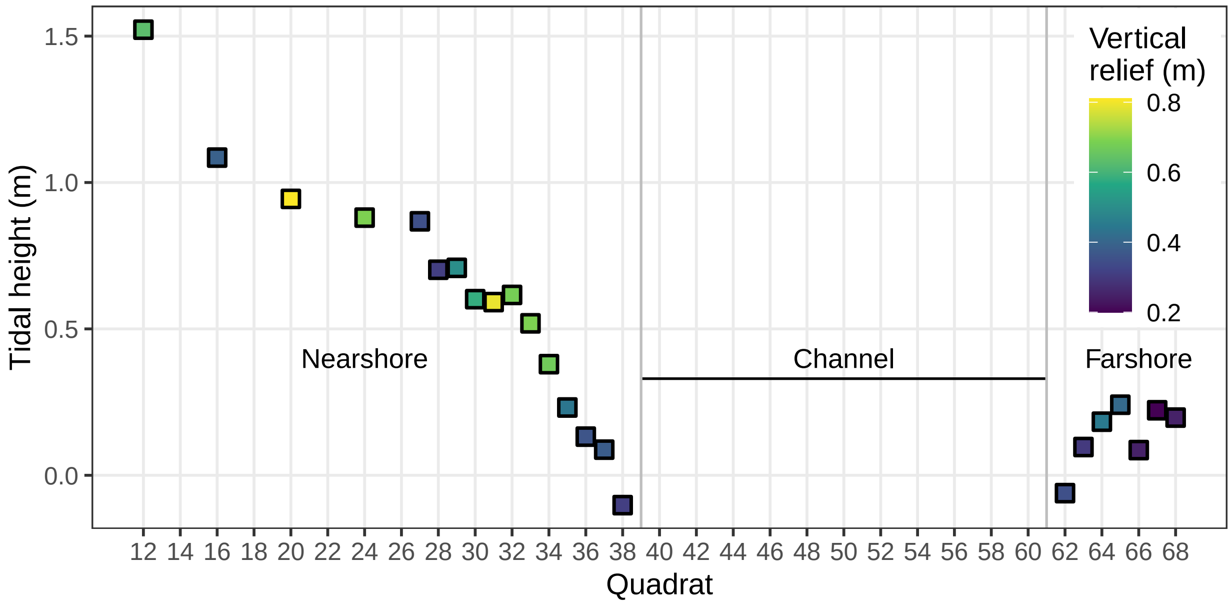


Figure S1. Visualization of the relative positions, tidal heights (m relative to mean lower low water), and vertical relief (m) of quadrats along Hewatt's transect that have been resurveyed since 1931. Quadrat 1 represents the start of Hewatt's transect at the uppermost limits of the rocky shore, and the transect runs perpendicular to the shoreline. Nearshore quadrats are separated from farshore quadrats by a channel at minus tides. Quadrats 27-68 are 'core' quadrats that were resurveyed from 1993 to 2023; quadrats 12-24 are 'extra' quadrats that were resurveyed from 2020 to 2023. Vertical relief was measured as the maximum vertical distance within a quadrat.

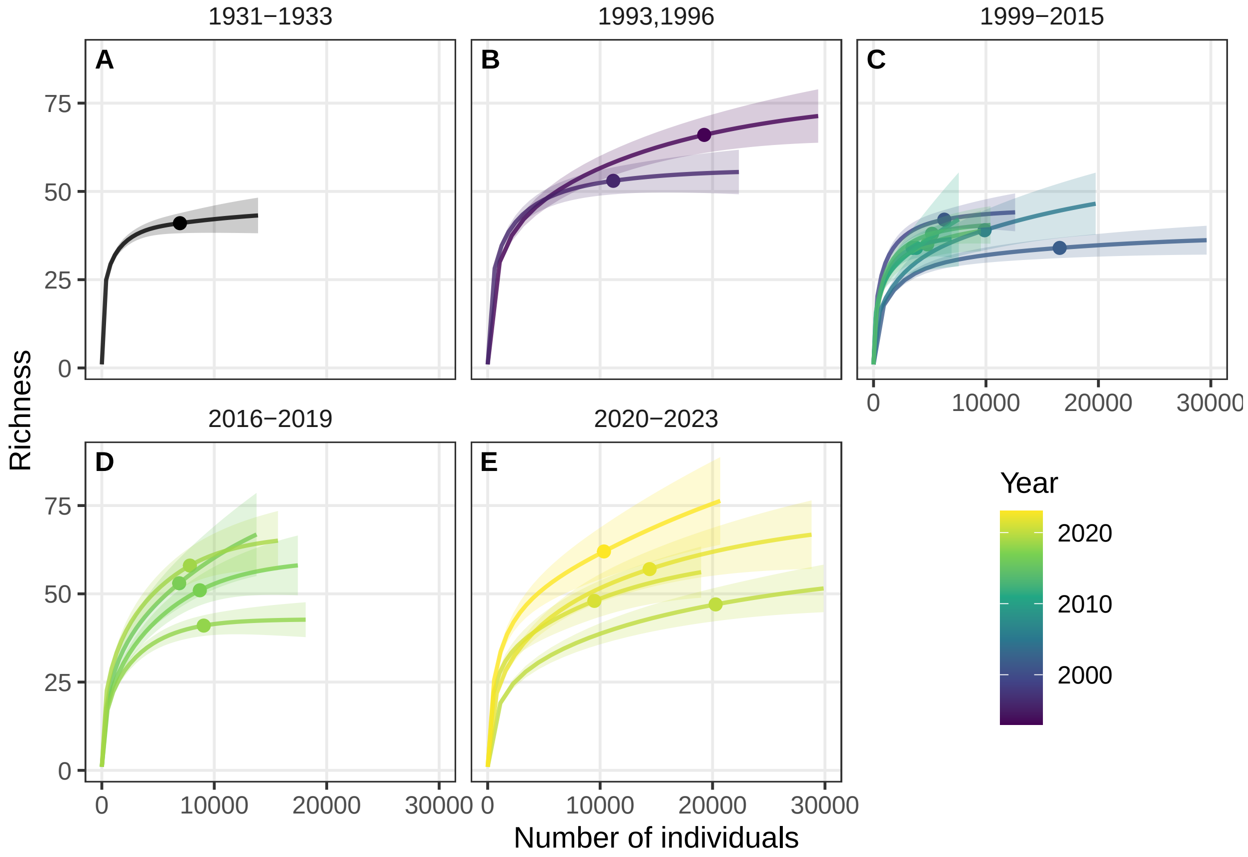


Figure S2. Species-accumulation curves from the repeated sampling conducted on Hewatt's transect during five investigator eras (1931-1933; 1993,1996; 1999-2015; 2016-2019; 2020-2023). The observed richness is plotted as a point; lines before the point represent rarefaction and lines beyond the point represent extrapolation. Shaded areas represent 95% confidence intervals.

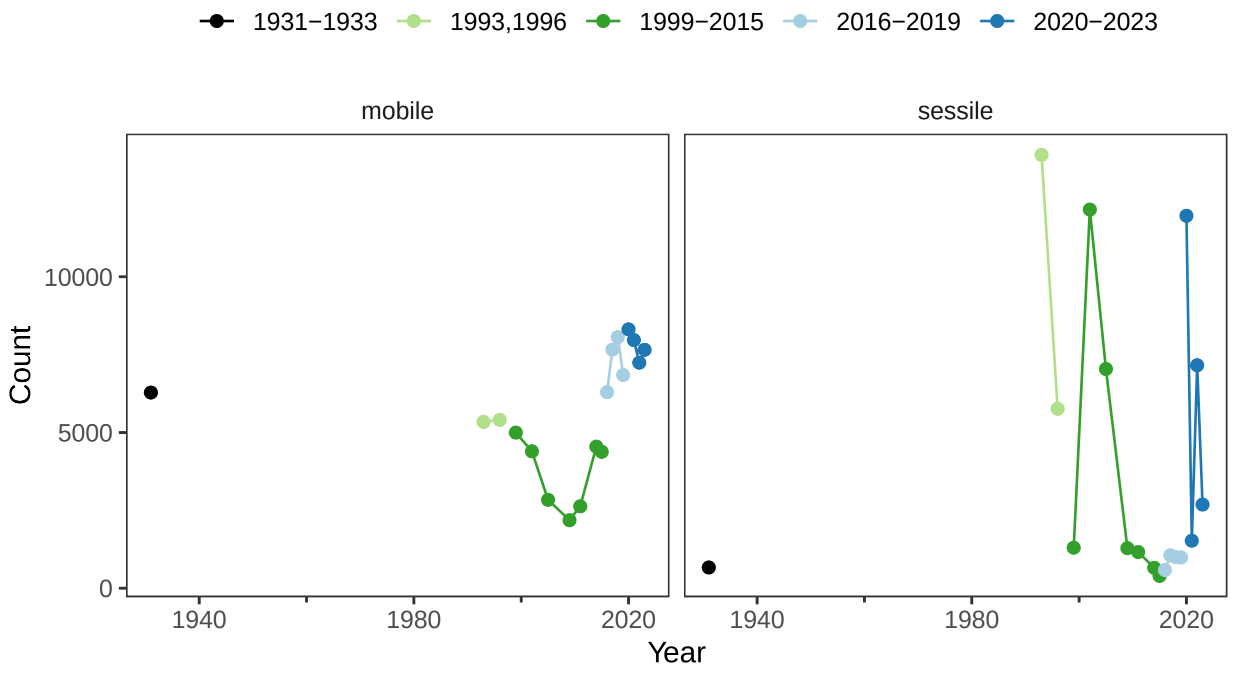


Figure S3. Total number of individuals from the repeated sampling conducted on Hewatt's transect, separated by motility (mobile vs sessile animals).

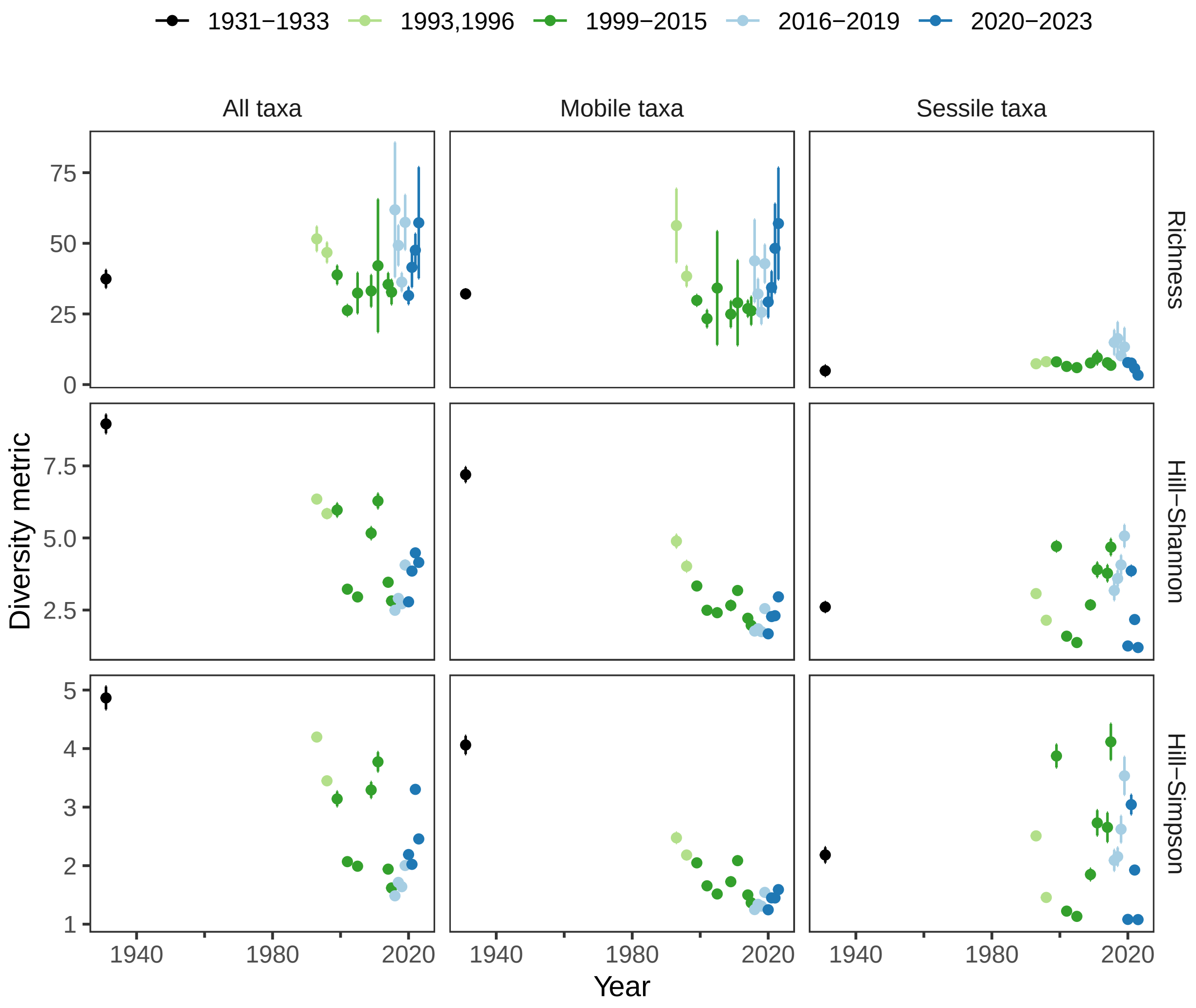


Figure S4. Temporal trends in site-scale alpha diversity metrics for the animal community on Hewatt's transect, separated by all taxa, mobile taxa, and sessile taxa. Diversity metrics were estimated using a coverage-based estimator (estimate ± 95% CI). Note that panels for all taxa are duplicated from Fig. 2D-F from the main text.
